# The Notch-Pdgfrβ axis suppresses brown adipocyte progenitor differentiation in early postnatal mice

**DOI:** 10.1101/2023.05.24.541839

**Authors:** Zuoxiao Shi, Shaolei Xiong, Ruoci Hu, Zilai Wang, Jooman Park, Yanyu Qian, Jaden Wang, Pratibha Bhalla, Nipun Velupally, Qing Song, Zhenyuan Song, Brian T. Layden, Yuwei Jiang

## Abstract

De novo brown adipogenesis holds potential in combating the epidemics of obesity and diabetes. However, the identity of brown adipocyte progenitor cells (APCs) and their regulation have not been extensively studied. Here through *in vivo* lineage tracing, we observed that PDGFRβ+ pericytes give rise to developmental brown adipocytes, but not to those in adult homeostasis. In contrast, TBX18+ pericytes contribute to brown adipogenesis throughout both developmental and adult stages, though in a depot-specific manner. Mechanistically, Notch inhibition in PDGFRβ+ pericytes promotes brown adipogenesis through the downregulation of PDGFRβ. Furthermore, inhibition of Notch signaling in PDGFRβ+ pericytes mitigates HFHS (high-fat, high-sucrose) induced glucose and metabolic impairment in both developmental and adult stages. Collectively, these findings show that the Notch/PDGFRβ axis negatively regulates developmental brown adipogenesis, and its repression promotes brown adipose tissue expansion and improves metabolic health.

**Highlights:**

- PDGFRβ+ pericytes act as an essential developmental brown APC.
- TBX18+ pericytes contribute to brown adipogenesis in a depot-specific manner.
- Inhibiting Notch-Pdgfrβ axis promotes brown APC adipogenesis.
- Enhanced postnatal brown adipogenesis improves metabolic health in adult stage.

## INTRODUCTION

Obesity has become a pressing health issue with its rising prevalence in both children and adults (Afshin et al., 2017). With obesity traditionally defined as the accumulation of excess fat, dysfunction in adipose tissue inevitably lies at the center of the pathogenic process of obesity and its related metabolic complications (Stefan et al., 2019). There are three types of adipocytes: white adipocytes, brown adipocytes, and, more recently identified, “brite” or beige adipocytes. White adipocytes primarily store excess energy, while beige and brown adipocytes, also known as thermogenic adipocytes, dissipate energy as heat to help maintain optimal temperatures for proper metabolic functions in organs.

Recent studies have provided mounting evidence that the activation or expansion of brown adipose tissue (BAT) holds potential in combating obesity and diabetes. For instance, BAT transplantation has been shown to promote glucose metabolism, reduce fat mass, and improve insulin resistance and liver steatosis in diabetic mice (Liu et al., 2015; Min et al., 2016; Zhang et al., 2021). Additionally, BAT has been demonstrated to have cardioprotective effects in several studies that employed mouse models with BAT dysfunction through either ablation of uncoupling protein 1 (UCP1)+ brown adipocytes or deletion of the UCP1 protein (Cittadini et al., 1999; Thoonen et al., 2015). In a more recent study, brown adipocytes with activated thermogenic metabolism were found to inhibit tumor growth by competing with cancer cells for glucose uptake, further emphasizing the therapeutic potential of brown adipocytes (Seki et al., 2022). Human studies have also provided evidence that presence of BAT is inversely associated with body mass index, and incidences of type II diabetes (T2D), hypertension and cardiovascular diseases, and others (Adachi et al., 2022; Becher et al., 2021; Betz and Enerback, 2011; Cai et al., 2019; Tam et al., 2012). These findings suggest its potential role in the development of obesity and its possible therapeutic significance. Indeed, the activation of BAT through cold exposure in obese humans can accelerate lipid metabolism, offering additional therapeutic benefits (Chondronikola et al., 2016).

Despite these findings, our understanding of BAT development and the molecular mechanisms regulating its formation remain relatively limited. BAT development begins during embryogenesis. In humans, BAT development ceases after infancy, whereas in rodents, BAT continues to develop, albeit at a slower pace, after birth (Wang et al., 2020). Numerous studies using Cre reporter mice have been conducted to understand BAT origin. Pax3+Pax7+Myf5+ cells have been found to give rise to interscapular BAT (iBAT) but not white adipocytes, suggesting a distinct lineage for brown and white adipocytes (Lepper and Fan, 2010; Shamsi et al., 2021). Interestingly, a separate study revealed that brown adipocytes in periaortic BAT (paBAT) arise from Wnt+ cells, not Myf5+ cells (Fu et al., 2019), suggesting the presence of depot-specific APCs. Moreover, in addition to iBAT, the largest and most extensively studied BAT depot in rodents, both humans and rodents have been found to possess at least four additional BAT depots (Mo et al., 2017; Zhang et al., 2018a). These include the supraclavicular BAT (scBAT), thoracic perivascular BAT (tPVAT), perirenal BAT (prBAT), and periaortic BAT (paBAT). More recently, the functions of these BAT depots are beginning to be appreciated. For example, scBAT in human infants seems to offer protection against excessive fat gain during the first six months of life (Entringer et al., 2017). Moreover, adipokines and extracellular vesicles from tPVAT have been implicated in pathological vascular remodeling associated with cardiovascular diseases (Barandier et al., 2005; Boucher et al., 2020; Huang et al., 2023; Li et al., 2019; Takaoka et al., 2009). The identification and significance of these new BAT depots underscore the complexity and heterogeneity of BAT, suggesting that research findings from one depot may not necessarily be applicable to others.

Mural cells, including pericytes associated with capillaries and venules, as well as vascular smooth muscle cells (VSMCs) linked with larger vessels like arteries and arterioles, have been identified as APCs for white adipose tissue (WAT) (Cawthorn et al., 2012; Gupta et al., 2012; Tang et al., 2008). These cells can differentiate into white adipocytes, thereby playing a crucial role in WAT homeostasis and expansion. Previously, our group proposed a multilineage model for white APCs based on developmental stages, suggesting that perivascular smooth muscle actin (SMA)+ cells act as adult-stage white APCs, while platelet-derived growth factor receptor alpha (PDGFRα)+ cells serve as developmental white APCs (Jiang et al., 2014; Shin et al., 2020). Others have shown that platelet-derived growth factor receptor beta (PDGFRβ)+ labeled pericytes can differentiate into mature white adipocytes under high-fat diet challenges and beige adipocytes under prolonged cold challenges, respectively (Vishvanath et al., 2015). Conversely, TBX18 labeled pericytes have not shown adipogenic potential (Guimarães-Camboa et al., 2017). However, the potential contribution of these cells to brown adipogenesis during the early postnatal stage and adult stage are largely unexplored, particularly in the smaller and more recently identified BAT depots.

Here, exploiting our two pericyte inducible reporter mice (PDGFRβ^RFP^ and TBX18^RFP^), we find that pericytes act as brown APCs in multiple BAT depots. Mechanistically, the adipogenic potential of these developmental APCs is repressed by the Notch signaling via its downstream target PDGFRβ. Deleting Rbpj, a major downstream effector of Notch signaling specifically in postnatal PDGFRβ+ pericytes, enhances their differentiation into brown adipocytes. Moreover, this enhanced postnatal brown adipogenesis alleviates high-fat, high-sucrose (HFHS) diet-induced glucose metabolism impairment and prevents HFHS diet-induced obesity in childhood and early adulthood. Thus, inhibiting Notch signaling in brown APCs may offer metabolic benefits for managing childhood and adult obesity, diabetes, and related complications.

## RESULTS

### Both PDGFRβ+ and TBX18+ pericytes contribute to brown adipogenesis during early postnatal stage

A recent study has highlighted that, in addition to cold challenges, mice during the early postnatal stage naturally develop beige adipocytes in the inguinal WAT (iWAT) (Chi et al., 2021). Concurrently, from postnatal 10 days (P10) to P30, BAT undergoes rapid expansion (Figure 1A). The tissue weight of iBAT increased by ∼150% (Figure 1A). To gain a comprehensive understanding of brown adipogenesis at this stage, we first confirmed their anatomic locations utilizing our inducible UCP1^RFP^ (Ucp1-Cre^ERT2^; Rosa26^RFP^) reporter mouse model with a single dose of Tamoxifen injection at P30, followed by analysis at P33 (Figure 1B). Whole-mount RFP imaging revealed the locations of RFP+ adipocytes in five BAT depots (iBAT, scBAT, tPVAT, prBAT, and paBAT) (Figure 1C-D). The presence of brown adipocytes was further confirmed through H&E staining, which showed adipocytes with a multilocular lipid droplet morphology, as well as through immunostaining of Perilipin (PLIN) and RFP (Figure 1E-G).

**Figure 1.**
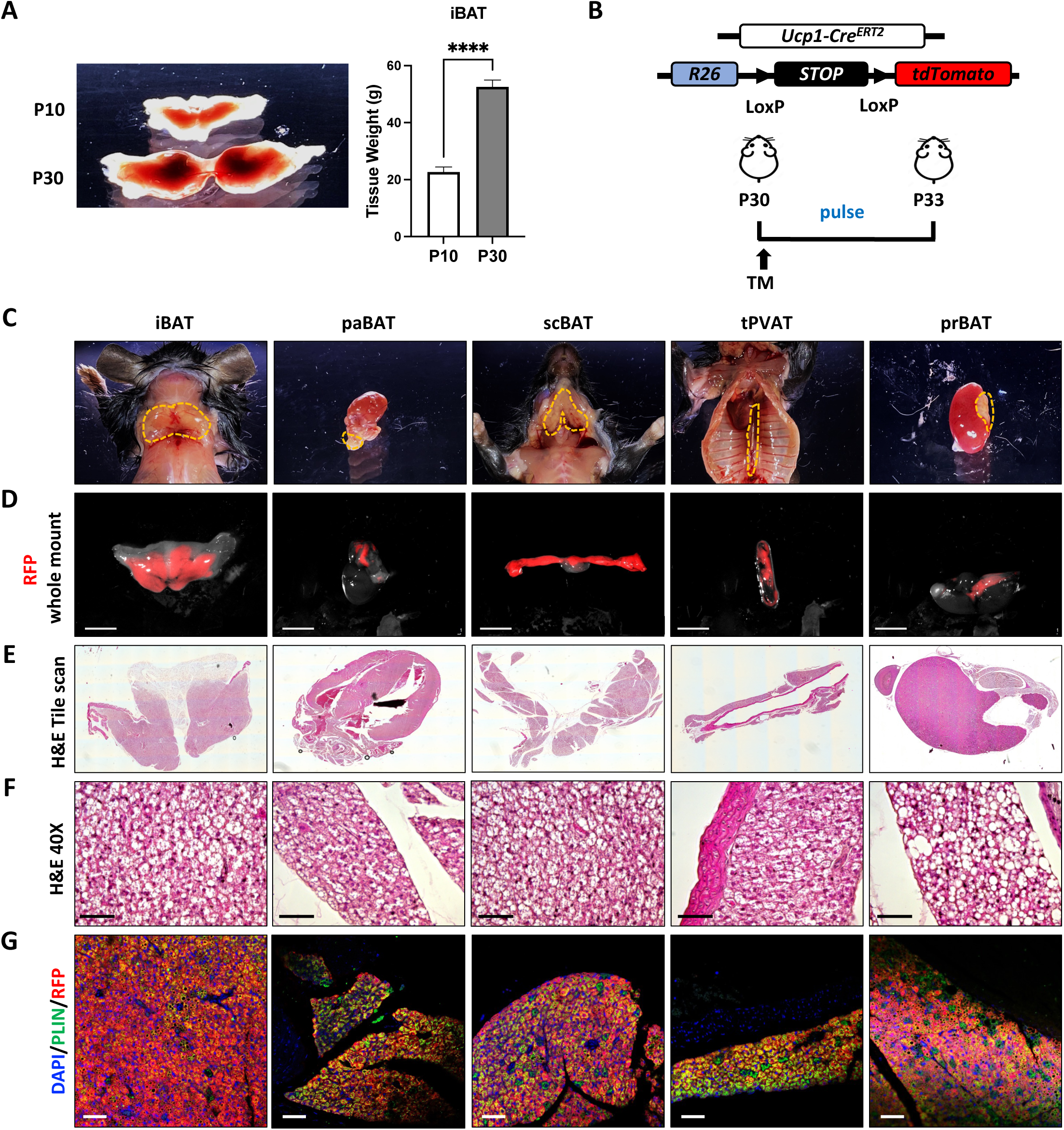
Anatomical locations of BAT depots and iBAT expansion during development. **(A)** Representative images of iBAT from P10 and P30 mice and tissue weights. **(B)** UCP1^RFP^ (Ucp1-Cre^ERT2^; Rosa26^RFP^) mouse model and experimental design. **(C)** Representative images of anatomical location of BAT depots. **(D)** Whole mount RFP images of BAT depots from (B). Scale bar, 1mm. **(E)** Tile scans of BAT depots in (D). **(E)** 40X representative images of brown adipocytes in (E). **(F)** IF staining of DAPI, PLIN and RFP of BAT depots in (D), Scale bar, 50 µm.

Pericytes have previously been found to give rise to mature white and beige adipocytes (Berry et al., 2016; Jiang et al., 2014; Vishvanath et al., 2015, 2016). To investigate the contribution of pericytes during BAT development, we crossed PDGFRβ-Cre^ERT2^ and TBX18-Cre^ERT2^ mice with Rosa26^RFP^ mice respectively (Figure 2A). Upon Tamoxifen injection, these mice undergo indelible RFP labeling of either PDGFRβ+ or TBX18+ cells. To confirm labeling of pericytes but not mature adipocytes, we first performed a pulse experiment by Tamoxifen injection at P10 and analyzed at P13. Whole mount images show presence of RFP+ cells in the BAT depots of interest. Furthermore, immunostaining of PLIN and RFP indicated RFP+ cells mainly localized to vesicular-like structure and RFP signal did not colocalize with PLIN, indicating exclusive labeling of RFP+ cells and brown adipocytes in all BAT depots in both reporter mice (Figure S1A-B). Next, to investigate pericytes contribution during early postnatal stage, we administered a single dose of Tamoxifen to P10 pups and performed fate mapping analysis at P30 (Figure 2B), using immunostaining with DAPI, PLIN and RFP. Whole mount images presented RFP+ tissues in the BAT depots of interest in both reporter mice, indicating P10 labeled pericytes are still present in the BAT depots (Figure 2C Upper Panel and Figure 2D Upper Panel). We found RFP+/PLIN+ cells in all five BAT depots of the PDGFRβ reporter mice (Figure 2C, Middle and Lower Panel). However, in the TBX18 reporter mice, these cells were only present in the scBAT, tPVAT, and prBAT depots (Figure 2D, Middle and Lower panel). Quantification of RFP+/PLIN+ pixels suggested varied contribution of PDGFRβ+ and TBX18+ cells in each BAT depot (Figure 2E). Together, our lineage tracing results suggest that PDGFRβ+ pericytes are universal brown APCs and TBX18+ pericytes are brown APCs of the scBAT, tPVAT and prBAT during early postnatal stage.

**Figure 2.**
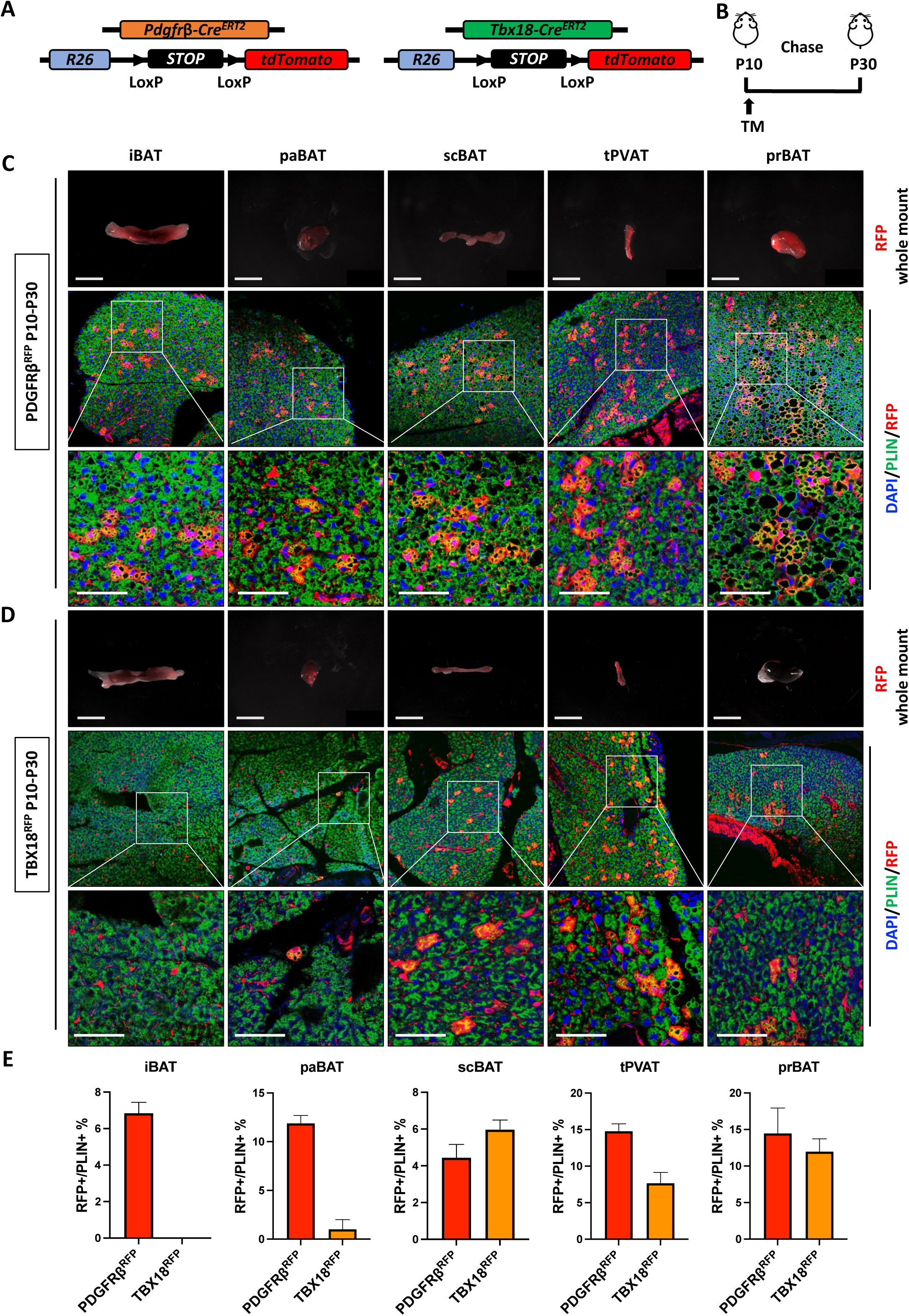
Pericytes contribute to brown adipogenesis during early postnatal stage. **(A)** PDGFRβ^RFP^ (Pdgfrβ-Cre^ERT2^; Rosa26^RFP^) and TBX18^RFP^ (Tbx18-Cre^ERT2^; Rosa26^RFP^) mouse models. **(B)** Experimental Design. TM injection at P10 and analysis at P30. **(C)** Upper panel: Whole mount RFP images of PDGFRβ^RFP^ BAT depots at P30. Scale bar, 1mm. Middle and lower panel: IF staining of DAPI (blue), PLIN (green) and RFP (red). Lower Panel: Zoomed-in images of middle panel. Scale Bar, 50 µm. **(D)** Upper panel: Whole mount RFP images of TBX18^RFP^ BAT depots at P30. Scale bar, 1mm. Middle and lower panel: IF staining of DAPI (blue), PLIN (green) and RFP (red). Lower Panel: Zoomed-in images of middle panel. Scale Bar, 50 µm. **(E)** Quantification of colocalization of PLIN and RFP of IF staining by pixels (n=3-4 images). Data are presented as means ± SEM.

### TBX18+ but not PDGFRβ+ pericytes give rise to brown adipocytes in the adult stage

To further investigate the adipogenic potential of pericytes across different BAT depots in the adult stage, we performed *in vivo* lineage tracing experiments in adult mice by Tamoxifen injection at P60 and analysis at P90. Whole mount images showed RFP+ signals in the BAT depots of interest from both reporter mice (Figure 3A-B, Upper panel), which indicates the presence of P60 labeled pericytes. To examine whether pericytes contribute to brown adipogenesis in adult stage, we performed immunostaining of DAPI, PLIN and RFP. In the immunostaining of BAT depots of PDGFRβ^RFP^ mice, no RFP+/PLIN+ cells were present in any of the depots, indicating that PDGFRβ+ pericytes do not contribute to brown adipogenesis during the adult stage under homeostatic conditions (Figure 3A, Middle and Lower panel). Conversely, in the BAT depots of TBX18^RFP^ mice, RFP+/PLIN+ cells were identified in the scBAT, tPVAT and prBAT, but not in the iBAT or the paBAT (Figure 3B, Middle and Lower panel). Collectively, these results indicate that while PDGFRβ+ pericytes do not contribute to brown adipogenesis during the adult stage under homeostatic conditions, TBX18+ pericytes are involved in the renewal of brown adipocytes specifically in the scBAT, tPVAT, and prBAT depots.

**Figure 3.**
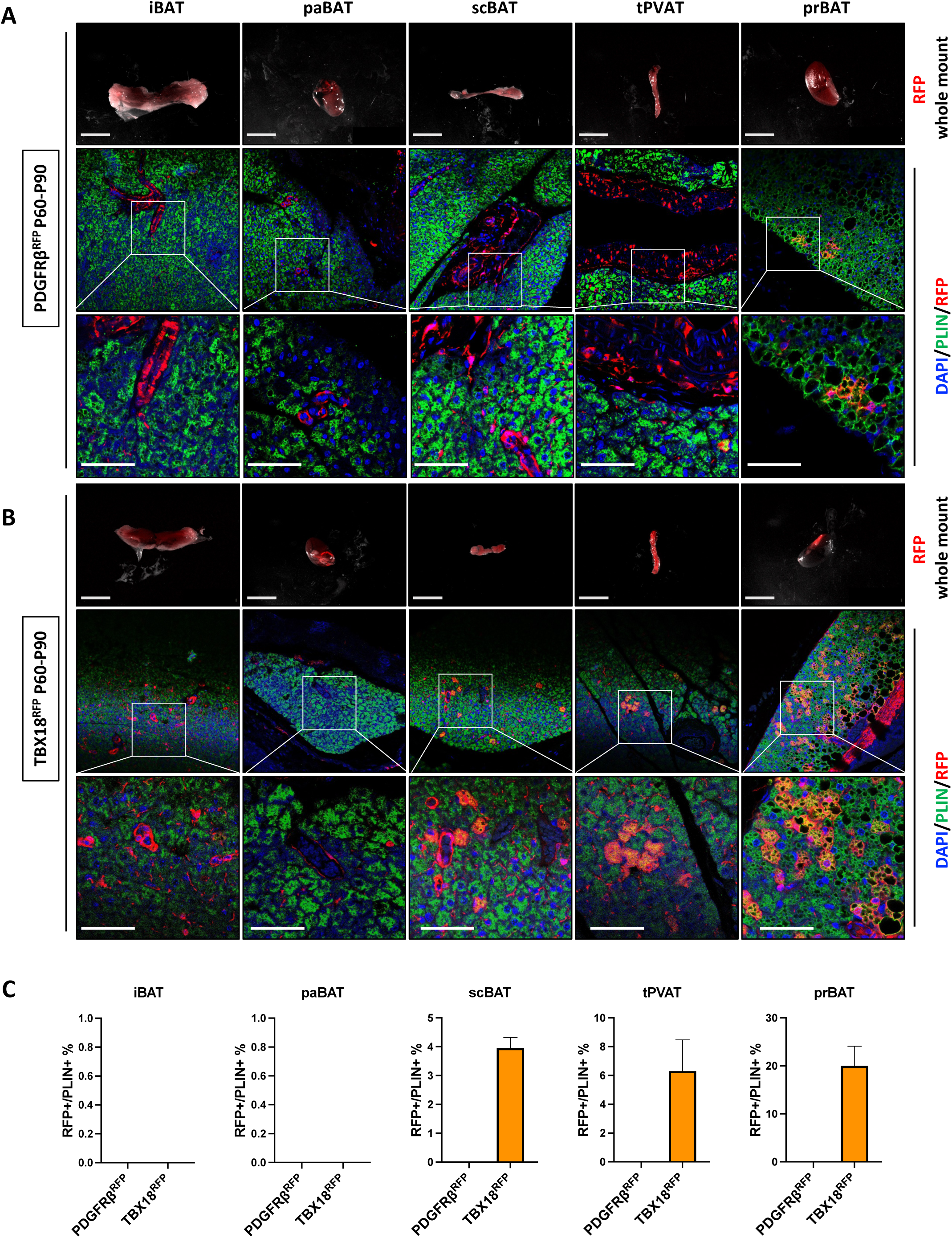
TBX18+ but not PDGFRβ+ pericytes give rise to brown adipocytes in the adult stage. **(A)** Upper panel: Whole mount RFP images of PDGFRβ^RFP^ BAT depots at P90. Scale bar, 1mm. Middle and lower panel: IF staining of DAPI (blue), PLIN (green) and RFP (red). Lower Panel: Zoomed-in images of middle panel. Scale Bar, 50 µm. **(B)** Upper panel: Whole mount RFP images of TBX18^RFP^ BAT depots at P90. Scale bar, 1mm. Middle and lower panel: IF staining of DAPI (blue), PLIN (green) and RFP (red). Lower Panel: Zoomed-in images of middle panel. Scale Bar, 50 µm. **(C)** Quantification of colocalization of PLIN and RFP of IF staining by pixels (n=3-4 images). Data are presented as means ± SEM.

### PDGFRβ+ pericytes commit to an adipocyte lineage during embryogenesis

To gain a more detailed understanding of pericyte contributions to BAT development in terms of timing, quantity, and anatomical location, we conducted a series of *in vivo* lineage tracing experiments on PDGFRβ^RFP^ mice using Tamoxifen injection at different time points (E14.5, P1, P5 or P10) and analyzed BAT depots with immunostaining at P30 (Figure 4A). We found that RFP+/PLIN+ cells were absent in the prBAT and paBAT in the E14.5-P30 group, indicating that PDGFRβ + pericytes do not contribute to these depots during this period. However, PDGFRβ+ pericytes contributed to brown adipocytes in the iBAT, scBAT and tPVAT starting at E14.5. In terms of quantity, PDGFRβ+ pericytes had the highest contribution to the iBAT and tPVAT when labeled at P1 and the highest contribution to scBAT, prBAT and paBAT when labeled at P5, suggesting a depot-specific variation in the contribution of PDGFRβ+ pericytes to brown adipogenesis (Figure 4B-C).

**Figure 4.**
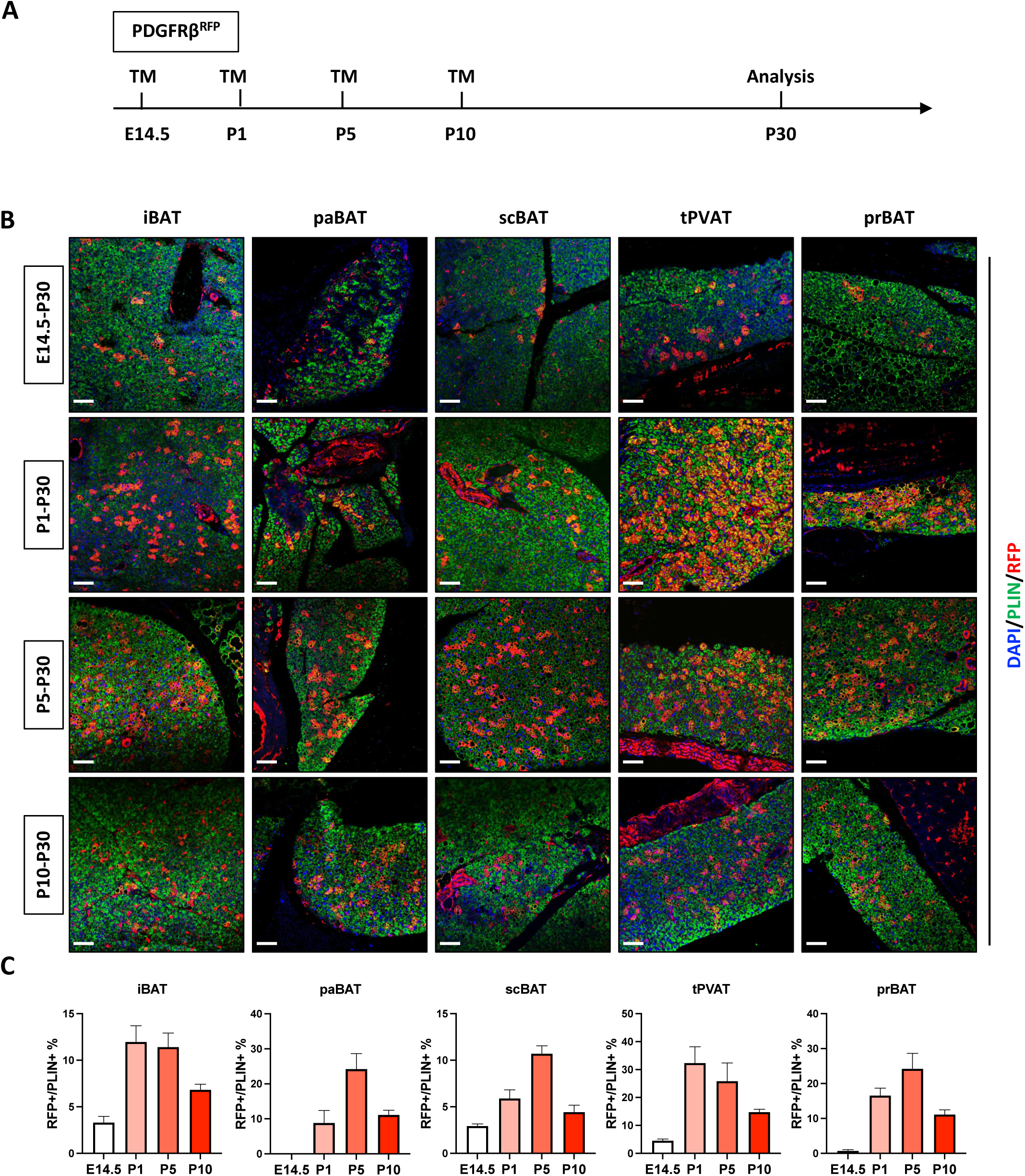
PDGFRβ+ pericytes acquire adipocyte fate during embryogenesis. **(A)** Experimental Design. TM injection at E14.5, or P1, or P5 or P10 and analysis at P30. **(B)** Representative images of IF staining of DAPI (blue), PLIN (green) and RFP (red). Scale bar, 50 µm. **(C)** Quantification of colocalization of PLIN and RFP of IF staining by pixels (n=3-4 images). Data are presented as means ± SEM.

Parallelly, similar experiments were also conducted in the TBX18^RFP^ mice with Tamoxifen injection at P1, P5 and P10 and subsequent analysis with immunostaining at P30. We found that RFP+ brown adipocytes were absent in P1, P5 and P10 iBAT (Figure S2A-B), suggesting that TBX18+ pericytes are not brown APCs of the iBAT during the postnatal stage. In addition, there were no RFP+ brown adipocytes in the paBAT with Tamoxifen injection at P10. However, P1 and P5 labeled TBX18+ pericytes gave rise to mature brown adipocytes at P30 (Figure S2A-B), indicating that TBX18+ pericytes contribute to paBAT development at early postnatal stage, which decreases with development. Similarly, TBX18+ pericyte contribution to the prBAT development starts as early as P1 and decreases over time (Figure S2A-B). Interestingly, TBX18+ pericyte contribution to scBAT and tPVAT starts as early as P1, with their contribution to these depots peaking at P5. Collectively, these results suggest that pericyte contribution to BAT development starts as early as embryogenesis and varies in terms of quantity based on developmental stage and location.

### PDGFRβ+ pericytes are essential for BAT development

To further investigate the importance of PDGFRβ+ cells in adipose tissue development, we constructed a mouse model PDGFRβ^DTA^ (PDGFRβ-Cre^ERT2^; Rosa26^DTA^; Rosa26^RFP^) that expresses diphtheria toxin subunit A (DTA) upon Tamoxifen injection, resulting in targeted ablation of PDGFRβ+ cells (Figure 5A). To assess its influence on BAT development, we injected Tamoxifen at P10 and analyzed the resulting metabolic phenotypes at P30 (Figure 5B). Our results showed that PDGFRβ^DTA^ mice had a small but significant reduction in body weight (Figure 5C) and in the tissue weights of iWAT, gonadal WAT (gWAT), and iBAT compared to the control PDGFRβ^RFP^ mice (Figure 5D). These findings indicate that PDGFRβ+ cells play an essential role in adipose tissue development. Meanwhile, the tissue weights of other organs, such as muscle, kidney, heart, spleen, liver, and pancreas remain similar between the two groups (Figure 5E), suggesting a specific role of PDGFRβ+ pericytes in the development of adipose tissues. In addition, fed blood glucose was not affected by the reduction in adipose tissue weights (Figure 5F). RFP immunostaining showed almost complete elimination of RFP+ brown adipocytes in the iBAT, scBAT, tPVAT and paBAT and a reduction in RFP+ brown adipocytes in the prBAT (Figure 5G-H). Of note, residual RFP+ pericytes were still observable in BAT depots of PDGFRβ^DTA^ mice, indicating incomplete ablation of PDGFRβ+ cells. To further confirm our results, we performed *in vitro* adipogenic assay on stromal vascular fraction (SVF) cells isolated from P10 Tamoxifen injected PDGFRβ^RFP^ and PDGFRβ^DTA^ mice at P13. Consistent with *in vivo* data, the percentage of RFP+ adipocytes in the SVF cells of all three BAT depots exhibited a significant reduction (Figure 5I). Collectively, our results suggest that PDGFRβ+ pericytes play a crucial role in BAT development.

**Figure 5.**
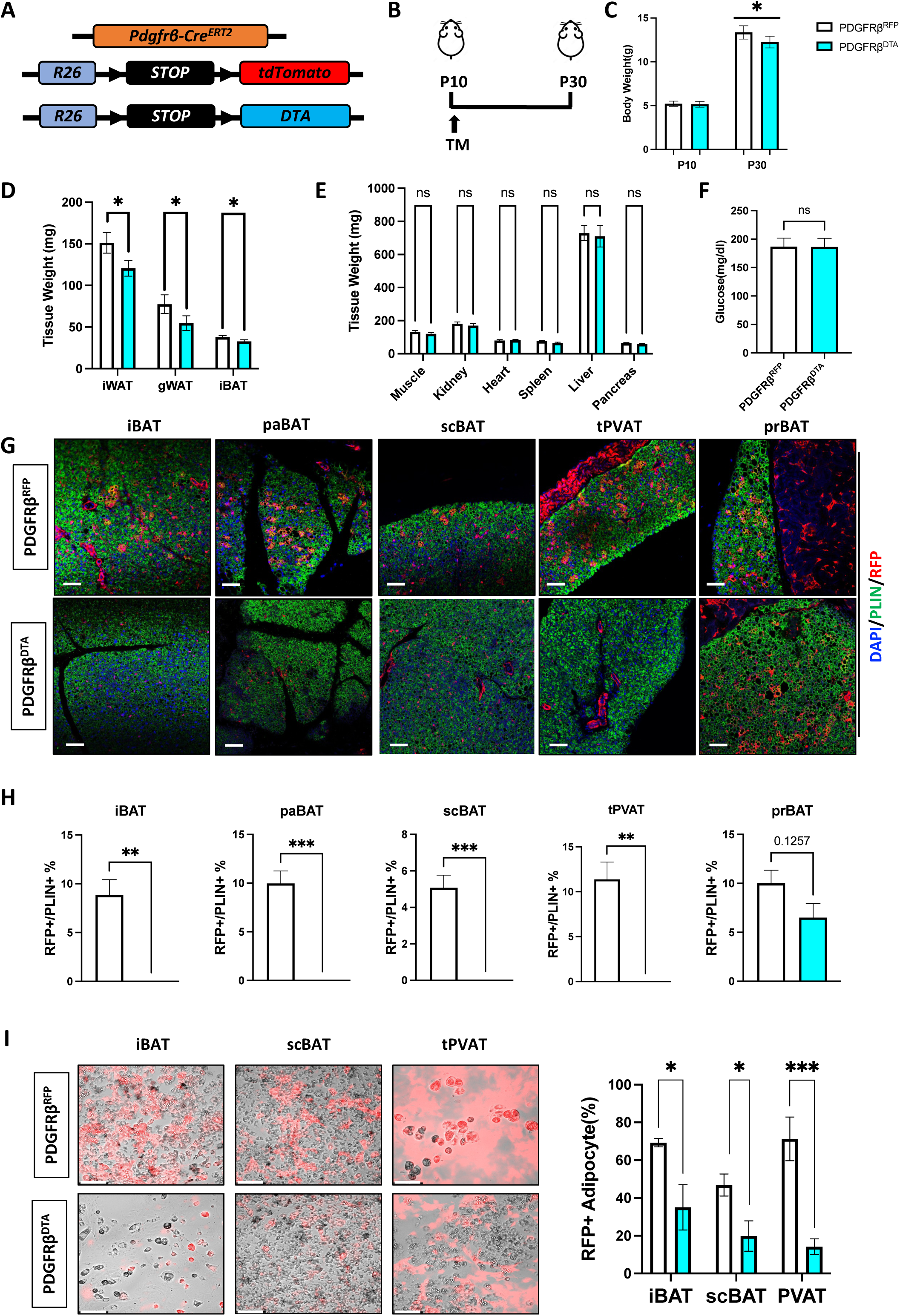
PDGFRβ+ pericytes are essential for brown adipose tissue development. **(A)** PDGFRβ^DTA^ (Pdgfrβ-Cre^ERT2^; Rosa26^DTA^) mouse model. **(B)** Experimental Design. TM injection at P10 and analyzed at P30. **(C)** Body weight (n=8-10). *p< 0.05. Data are presented as means ± SEM. **(D)** Adipose tissue weight (n=8-10). *p< 0.05. Data are presented as means ± SEM. **(E)** Other organ tissue weight (n=8-10). Data are presented as means ± SEM. **(F)** Fed blood glucose level. Data are presented as means ± SEM. **(G)** Representative images of IF staining of DAPI (blue), PLIN (green) and RFP (red). Scale bar, 50 µm. **(H)** Quantification of colocalization of PLIN and RFP of IF staining by pixels (n=3-4 images). Data are presented as means ± SEM. **(I)** DIC/RFP representative images of differentiated SVF cells from BAT depots. Quantification of RFP+ adipocytes% in PDGFRβ^RFP^ and PDGFRβ^DTA^ mice. *p< 0.05, ***p< 0.001. Data are presented as means ± SEM.

### Inhibiting Notch signaling in PDGFRβ+ pericytes promotes brown adipogenesis

Previous studies have shown that Notch signaling can drive adipocyte dedifferentiation and tumorigenic transformation in mice (Bi et al., 2016). Conversely, suppressing Notch signaling in white adipocytes can induce adipocyte browning through regulating UCP1 expression (Bi et al., 2014). We reasoned that Notch signaling may regulate differentiation of PDGFRβ+ brown APCs, thereby impacting brown adipogenesis. To investigate this, we examined whether Notch signaling genes were enriched in PDGFRβ+ brown APCs. Through gene expression analysis of sorted RFP+ and RFP-cells from P10 Tamoxifen-injected PDGFRβ^RFP^ mice, we validated the enrichment of *Pdgfrβ* and *Rfp* expression in the RFP+ population (Figure 6A). Furthermore, we noticed an increased presence of *Notch2* and *Notch3* expression in the RFP+ population, suggesting a potential role for Notch signaling in PDGFRβ+ brown APCs (Figure 6A).

**Figure 6.**
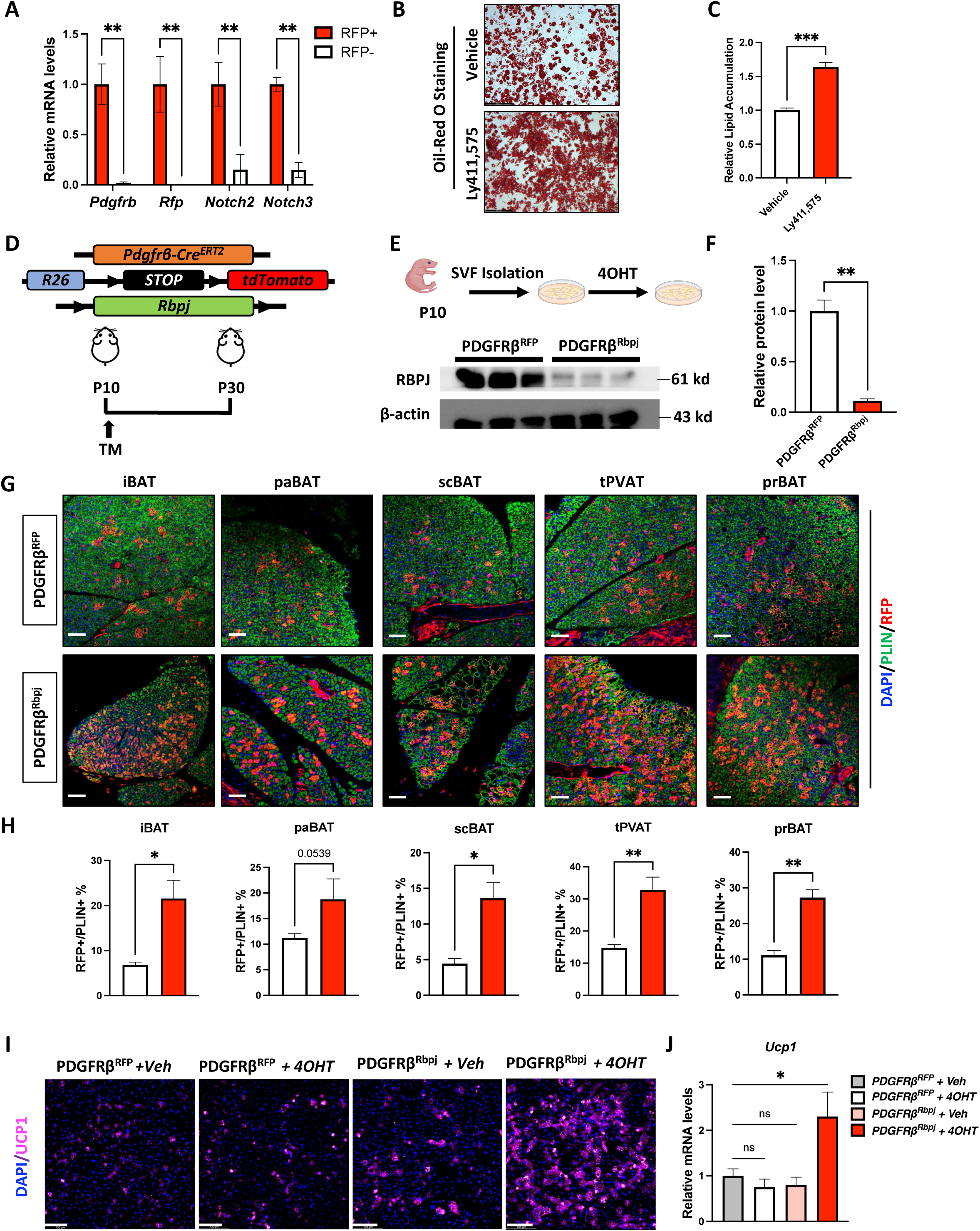
Inhibiting notch signaling in PDGFRβ+ pericytes promotes brown adipogenesis. **(A)** qRT-PCR analysis of RFP+ and RFP-cell populations sorted SVF cells from PDGFRβ^RFP^ mice. (n=3 per group). *p<0.05, **p<0.01. Data are presented as means ± SEM. **(B)** Representative images of Oil-Red O staining of SVF cells pretreated with vehicle or Ly411,575 before differentiation. **(C)** Quantification of relative Oil Red O accumulation from (B). ***p<0.001. Data are presented as means ± SEM. **(D)** PDGFRβ^Rbpj^ (Pdgfrβ-Cre^ERT2^; Rosa26^RFP^; Rbpj^fl/fl^) mouse model. **(E)** Upper panel: experimental design. Lower panel: western blot of RBPJ and β-actin from SVF cells of PDGFRβ^RFP^ and PDGFRβ^Rbpj^ mice. **(F)** Quantification of western blot from (E). **p<0.01. Data are presented as means ± SEM. **(G)** Representative images of IF staining of DAPI (blue), PLIN (green) and RFP (red). Scale bar, 50 µm. **(H)** Quantification of colocalization of PLIN and RFP of IF staining by pixels (n=3-4 images). *p<0.05, **p<0.01. Data are presented as means ± SEM. **(I)** Representative image of IF staining of DAPI (blue) and UCP1(purple). Scale bar, 50 µm **(J)** qRT-PCR analysis of Ucp1. (n=3 per group). *p<0.05. Data are presented as means ± SEM.

To investigate the role of Notch signaling in APCs during development, we pre-treated SVF cells with Ly411,575 (100 nM), a gamma-secretase inhibitor, until the cells reach 85% confluency. We then differentiated the cells into mature adipocytes with insulin. Oil Red O staining showed a significant increase in adipogenesis and lipid accumulation, suggesting a negative regulatory role of Notch signaling in adipogenesis (Figure 6B-C). To specifically investigate the effects of Notch inhibition in PDGFRβ+ pericytes, we generated a PDGFRβ^Rbpj^ mouse model (PDGFRβ-Cre^ERT2^; Rbpj^fl/fl^; Rosa26^RFP^) (Figure 6D, Upper panel). In this model, upon Tamoxifen injection, Rbpj, the major effector of Notch signaling, would be knocked out. To verify Rbpj knockout efficiency, we cultured SVF cells isolated from PDGFRβ^RFP^ and PDGFRβ^Rbpj^ mice and treated them with 4-OHT, the active metabolite of Tamoxifen to trigger Cre translocation and genetic deletion. Western blot analysis showed ∼90% reduction in RBPJ in protein level (Figure 6E-F). To examine the effect of RBPJ knockout on BAT development, we injected Tamoxifen at P10 and analyzed the BAT depots at P30 (Figure 6D, Lower panel). PDGFRβ^RBPJ^ mice showed an increased number of RFP+ adipocytes in all BAT depots of interest, compared to the control groups (Figure 6G-H). To further test if Notch inhibition affects brown adipogenesis, we treated SVF cells from PDGFRβ^RFP^ and PDGFRβ^Rbpj^ mice with either vehicle or 4OHT to induce RBPJ deletion, followed by a brown adipogenesis differentiation protocol. Immunostaining of DAPI/UCP1 and qRT-PCR analysis of *Ucp1* level showed only increased UCP1 staining and expression in PDGFRβ^Rbpj^ + 4OHT treatment group (Figure 6I-J), confirming the inhibitory role of Notch in brown adipogenesis. Taken together, our data indicate that Notch signaling inhibits the differentiation of PDGFRβ+ pericytes into brown adipocytes.

### Notch inhibition promotes brown adipogenesis through Pdgfrβ downregulation

Notch signaling exerts its effects through translocation of the intracellular domain (NICD) into the nucleus and direct binding with targeted genes with other transcription factors (Kangsamaksin et al., 2014). One of the potential downstream targets of Notch signaling is PDGFRβ, whose first intron region has been shown to be bound by Rbpj upon Notch activation (Castel et al., 2013). Two other studies have also shown that Notch activation promotes PDGFRβ expression and signaling when overexpressing NICD or deleting its downstream effectors (Jin et al., 2008; Nadeem et al., 2020). In conditions of obesity and in models with pharmacologically induced inhibition of adipogenesis, upregulation of PDGFRβ ligands has been observed (Cheng et al., 2021; Shao et al., 2021a; Zhang et al., 2018b). This suggests that PDGFRβ could potentially serve as a downstream effector in the Notch inhibition pathway, thereby promoting adipogenesis. Consistent with this hypothesis, we noted a significant downregulation of PDGFRβ in SVF cells isolated from PDGFRβ^Rbpj^ mice (Figure S3A-B). Additionally, we noticed a reduction in adipogenic potential of SVF cells from adult P90 mice compared to P10 pups (Figure S3C-D). This was coupled with an enhancement of both Notch and PDGFRβ signaling gene expression (Figure S3E). To verify the inhibitory effect of PDGFRβ signaling in adipogenesis, we pretreated P10 SVF cells with PDGFRβ activator PDGF-DD (25ng/ml) or PDGFRβ inhibitors, APB5 (PDGFRβ antibody) (0.5 ug/ml) and Sunatinib (PDGFRβ inhibitor) (100nM) followed by insulin for adipocyte differentiation. PDGFRβ inhibitor and antibody pretreatment increased adipogenesis whereas PDGF-DD pretreatment inhibited adipogenesis (Figure S3F-G). To test if PDGFRβ signaling is downstream of Notch signaling, we pretreated P10 SVF cells with either a Notch inhibitor (Ly411,575) alone, or a combination of Notch inhibitor and PDGFRβ activator (PDGF-DD). The promotion of adipogenesis observed with Notch inhibitor treatment was attenuated by PDGF-DD, as indicated by Oil Red O staining and analysis of adipogenic gene expression (Figure S3H-J). To further understand how PDGFRβ signaling affect the brown adipogenic process, we performed a time course treatment of PDGF-DD during the differentiation process. As a result, the gene expression level of *Pparγ2*, a master regulator of adipogenesis, was shown to be downregulated by PDGF-DD treatment (Figure S3K). Taken together, our results suggest that the promotion of adipogenesis via Notch inhibition might potentially be achieved through the downregulation of PDGFRβ expression and its subsequent impact on Pparγ2 signaling.

### Notch inhibition in PDGFRβ+ cells prevents HFHS diet-induced impairment of glucose metabolism during childhood

The consumption of a high-fat, high-sucrose (HFHS) diet is known to contribute to the development of obesity and diabetes. To investigate whether the enhanced brown adipogenesis in PDGFRβ^Rbpj^ mice has beneficial effects in preventing childhood obesity and its associated metabolic disorders, we challenged P10 Tamoxifen-injected PDGFRβ^RFP^ and PDGFRβ^Rbpj^ mice with an HFHS diet until P30 for analysis (Figure 7A). We found that PDGFRβ^Rbpj^ mice had a significantly reduced body weight gain from HFHS challenge compared to its chow counterpart (Figure 7B and Figure S4A), which was not due to food intake (Figure S4B). Fat or lean mass percentages did not differ between the two groups (Figure S4C-D). In agreement with the lower increase in body weight, the PDGFRβ^Rbpj^ mice had an improvement of glucose metabolism, indicated by significantly lower fed blood glucose level (Figure S4E) and improved glucose tolerance compared with PDGFRβ^RFP^ mice (Figure 7C-D). Additionally, we found PDGFRβ^Rbpj^ had significantly lower triglyceride level in the blood serum and liver (Figure S4F). Consistently, we found PDGFRβ^Rbpj^ mice exhibited less microvesicular steatosis compared to control group, as revealed by liver H&E staining (Figure 7E). Although there was no significant difference in tissue weights of the iWAT and gWAT (Figure S4G), the iBAT tissue weight was significantly higher in the PDGFRβ^Rbpj^ mice (Figure 7F), suggesting potentially increased thermogenic activity. Indeed, upon acute cold challenge, the PDGFRβ^Rbpj^ mice were more capable of maintaining their rectal body temperature compared to the PDGFRβ^RFP^ mice (Figure 7G). H&E staining did not show observable difference in brown adipocyte morphology in the five BAT depots (Figure S4H). However, white adipocyte size analysis showed an increased percentage of smaller adipocytes in the PDGFRβ^Rbpj^ mice (Figure S4I), suggesting a better metabolic health. And consistent with the previous findings, immunostaining of BAT depots in PDGFRβ^Rbpj^ mice presented with more RFP+ brown adipocytes (Figure 7H-I). Interestingly, despite no significant changes observed in thermogenic gene expression in iBAT (Figure S4J), our gene expression analysis revealed a significant upregulation of several rate-limiting enzymes involved in lipid and glucose metabolism (Figure 7J-K). This finding suggests that the improved metabolic health observed in our study may be attributed, at least in part, to enhanced lipid and glucose processing pathways rather than alterations in thermogenesis. Collectively, our data suggest that promoted brown adipogenesis through Rbpj knockout in PDGFRβ+ pericytes under an HFHS diet not only prevented body weight gain but also significantly improved overall metabolic health.

**Figure 7.**
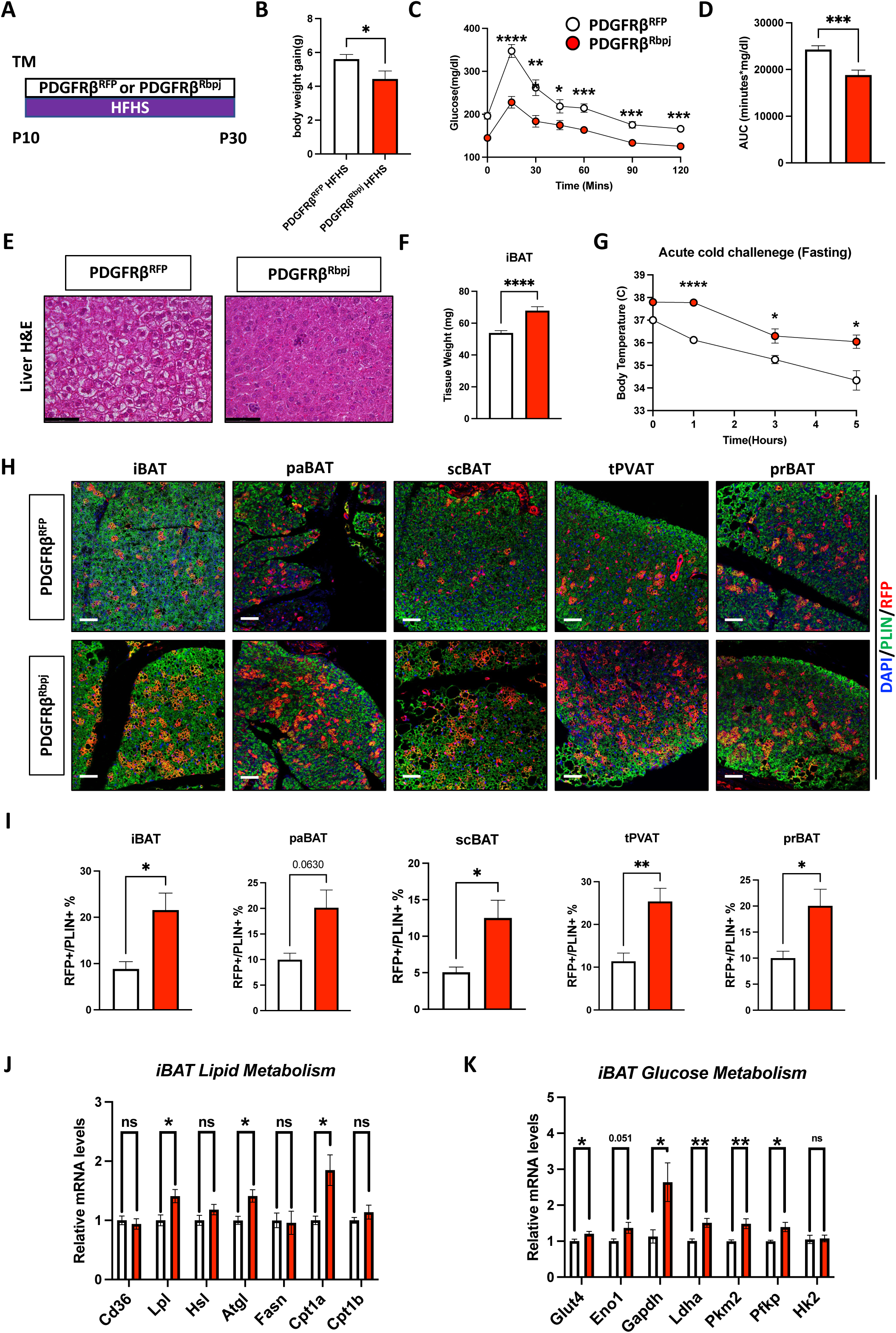
Notch inhibition in PDGFRβ+ cells prevents HFHS diet-induced impairment of glucose metabolism during development. **(A)** Experimental Design. TM injection at P10 followed by HFHS feeding for 20 days and analysis at P30. **(B)** Body weight gain compared to chow counterparts (n>10 mice per group). *p<0.05. Data are presented as means ± SEM. **(C)** GTT Curve (n>10 mice per group). *p<0.05, **p<0.01, ***p<0.001, ****p<0.0001. Data are presented as means ± SEM. **(D)** GTT AUC quantification. ***p<0.001. Data are presented as means ± SEM. **(E)** H&E staining of Liver. Scale bar, 50 µm. **(F)** iBAT tissue weights (n>10 per group). ****p<0.0001. Data are presented as means ± SEM. **(G)** Rectal body temperature during cold challenge with fasting (n= 5, 4). *p<0.05, ****p<0.0001. Data are presented as means ± SEM. **(H)** Representative images of IF staining of DAPI (blue), PLIN (green) and RFP (red). Scale bar, 50 µm. **(I)** Quantification of colocalization of PLIN and RFP of IF staining by pixels (n=3-4 images). *p<0.05, **p<0.01. Data are presented as means ± SEM. **(J)** qRT-PCR analysis of lipid metabolism genes (n=5 per group). *p<0.05. Data are presented as means ± SEM. **(K)** qRT-PCR analysis of glucose metabolism genes (n=5 per group). *p<0.05, **p<0.01. Data are presented as means ± SEM.

### Increased brown adipogenesis during development prevents early adult obesity

To further explore whether the increased brown adipogenesis in PDGFRβ^Rbpj^ during the postnatal stage has metabolic benefits in early adult life, we challenged P10 Tamoxifen-injected PDGFRβ^RFP^ and PDGFRβ^Rbpj^ mice to a 30-day challenge with a HFHS diet starting at P30 (Figure 8A). At P30, PDGFRβ^Rbpj^ mice had a higher body weight compared to controls (Figure S5A). However, after 30 days on the HFHS diet, PDGFRβ^Rbpj^ mice gained significantly less body weight (Figure 8B). Consistently, despite no significant difference in food intake (Figure S5B), PDGFRβ^Rbpj^ mice had a significant reduction in body fat mass percentage and a significantly higher lean mass percentage (Figure 8C-D). Concurrently, glucose metabolism was also improved in PDGFRβ^Rbpj^ mice, assessed by reduced fed blood glucose levels and significantly improved glucose tolerance (Figure 8E-G). Furthermore, the enhanced metabolic fitness was further supported by a significant reduction in serum triglyceride levels and a decrease in liver cholesterol content in PDGFRβ^Rbpj^ mice (Figure S5C). In addition, PDGFRβ^Rbpj^ mice had significantly lowered tissue weights in iWAT and gWAT, but significantly higher iBAT weights, (Figure 8H). PDGFRβ^Rbpj^ mice also exhibited a shift in white adipocyte size distribution towards smaller adipocytes, suggesting hyperplasia over hypertrophy (Figure S5D-E). In addition, the liver of PDGFRβ^RFP^ mice displayed apparent lipid droplet accumulation, indicating steatosis, while the liver of PDGFRβ^Rbpj^ mice exhibited mild microvesicular steatosis (Figure 8I). There was not a significant difference in the percentage of RFP+ brown adipocytes in the BAT depots under study (Figure 8J). However, given the increased iBAT tissue weight, the total contribution of brown adipocytes from PDGFRβ+ cells was anticipated to be higher in the PDGFRβ^Rbpj^ mice. Increased iBAT weight did not result in increased thermogenic gene expression changes (Figure S5F). However, consistent with developmental stage HFHS diet treated mice, iBAT of PDGFRβ^Rbpj^ mice had significantly increased lipid and glucose metabolism gene expressions (Figure 8K-L). Consistently, H&E staining also showed less whitening of brown adipocytes in PDGFRβ^Rbpj^ compared to PDGFRβ^RFP^ mice, suggesting a potential improved function of iBATs (Figure S5G). Together, these findings suggest that promoting thermogenic adipogenesis during the developmental stage can confer long-lasting beneficial effects into early adult life, emphasizing its importance in promoting overall metabolic health throughout the lifespan.

**Figure 8.**
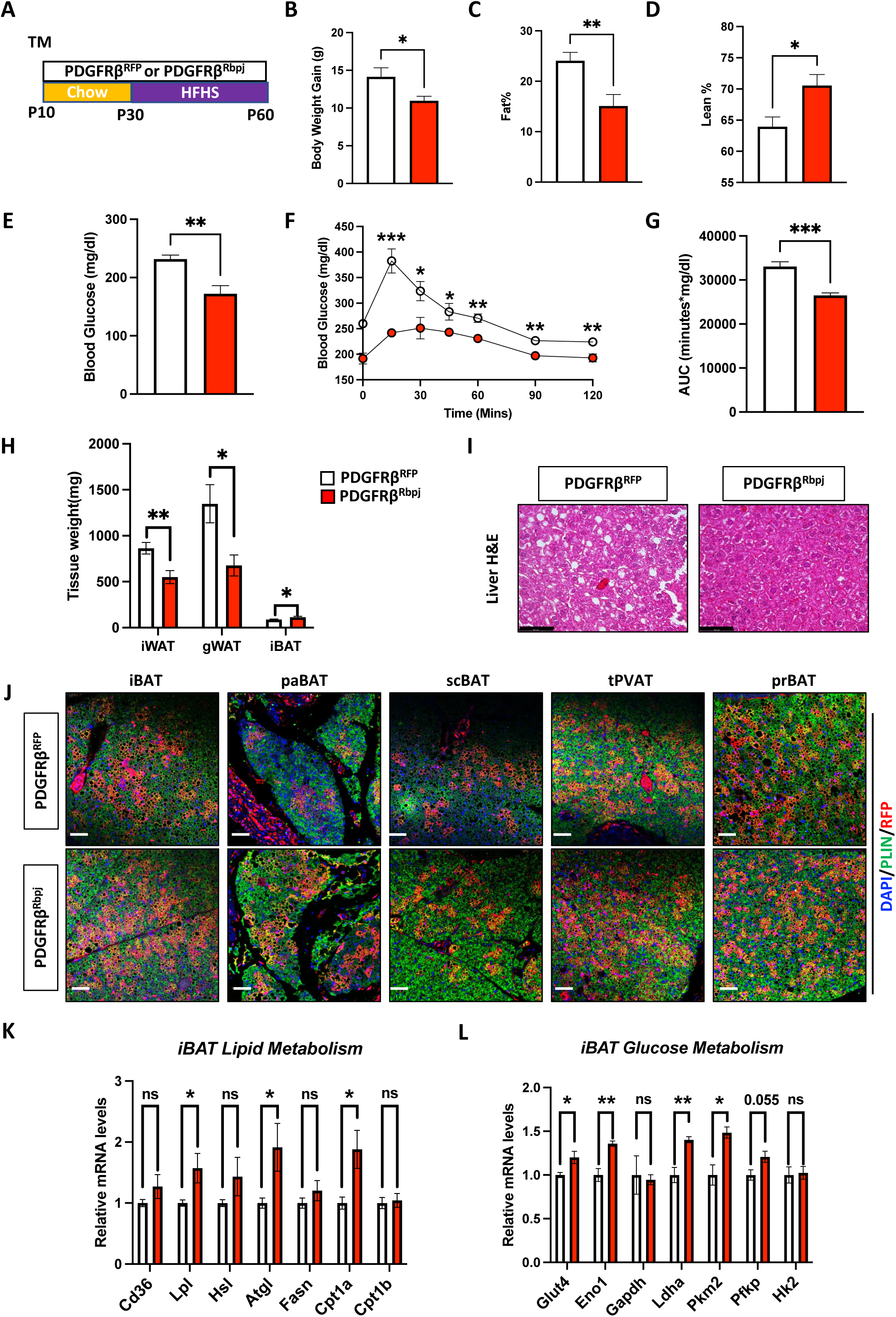
Increased brown adipogenesis during development prevents adult obesity. **(A)** Experimental Design. TM injection at P10 followed by HFHS feeding for 30 days starting at P30. **(B)** Body weight gain compared to P30 (n=6 mice per group). *p<0.05. Data are presented as means ± SEM. **(C)** Fat content at P60 (n=6 per group). **p<0.01. Data are presented as means ± SEM. **(D)** Lean content at P60 (n=6 per group). *p<0.05. Data are presented as means ± SEM. **(E)** Fed blood glucose level (n=6 per group). **p<0.01. Data are presented as means ± SEM. **(F)** GTT Curve (n=6 per group). *p<0.05, **p<0.01, ***p<0.001. Data are presented as means ± SEM. **(G)** GTT AUC quantification (n=6 per group). ***p<0.001. Data are presented as means ± SEM. **(H)** Adipose tissue weights at P60 (n=6 per group). *p<0.05, **p<0.01. Data are presented as means ± SEM. **(I)** Liver H&E staining. Scale bar, 50 µm. **(J)** Representative images of IF staining of DAPI (blue), PLIN (green) and RFP (red). Scale bar, 50 µm. **(K)** qRT-PCR analysis of lipid metabolism genes (n=5 per group). *p<0.05. Data are presented as means ± SEM. **(L)** qRT-PCR analysis of glucose metabolism genes (n=5 per group). *p<0.05, **p<0.01. Data are presented as means ± SEM.

## DISCUSSION

Extensive studies in the past have focused on identifying APCs in the adult stage for white and beige adipocytes, along with their regulatory mechanisms. In contrast, there has been less focus on the identification and regulatory mechanisms of brown APCs, especially during developmental stages. To fill this knowledge gap, we conducted lineage tracing studies to identify APCs for various BAT depots during both early postnatal and adult stages. BAT development occurs during late embryogenesis and continues into early postnatal stage. For instance, UCP1+ brown adipocytes begin to form in the iBAT from E16.5, and the paBAT starts developing from E18.5.(Angueira et al., 2021; Mayeuf-Louchart et al., 2019). Another study utilizing the AdipoChaser mouse model, the development of brown adipocytes in the iBAT starts as early as E10 and completes by E16, as no new brown adipocytes were observed to be labeled beyond this point (Song et al., 2020). However, our research findings suggest that pericytes give rise to mature brown adipocytes in multiple BAT depots during the postnatal phase. Specifically, PDGFRβ+ pericytes serve as a universal APCs for all BAT depots studied during developmental stage, while TBX18+ pericytes function as brown APCs for the tPVAT, scBAT, and prBAT. Interestingly, TBX18+ pericytes labelled at P10 do not contribute to brown adipogenesis in paBAT, whereas those labelled at the perinatal age of P1 do contribute to this process. This suggests that the contribution of pericytes to brown adipogenesis is contingent upon the developmental stage, and the timeframe for this difference could be as brief as 10 days, underscoring the temporal specificity of this developmental process. It is worth noting that the PDGFRβ-Cre reporter mouse model has been criticized for its early expression during embryogenesis, making it less suitable for lineage tracing (Guimarães-Camboa et al., 2017). Our results suggest that PDGFRβ+ pericytes labeled at E14.5 contribute minimally to brown adipogenesis compared to those labeled at P1 and P5. This suggests that the contribution of pericytes to brown adipogenesis may occur concurrently with organogenesis rather than preceding it.

Another interesting observation is that brown adipocytes continue to be generated during adulthood under homeostatic conditions. Typically, BAT is most abundant in newborns and declines with age (Shamsi et al., 2021). However, there is limited evidence regarding whether BAT continues to produce new brown adipocytes during adult homeostasis or during the physiological changes associated with aging. Our study provides evidence that, under homeostatic conditions, brown adipocytes continue to be produced in adulthood, with TBX18+ pericytes, but not PDGFRβ+ pericytes, serving as their APCs. Interestingly, this production is only observed in the scBAT, tPVAT and prBAT, and not in the iBAT or the paBAT. These findings align with previous research by Dr. Sylvia Evans’ group, indicating that TBX18+ pericytes do not function as tissue-resident multipotent progenitors in various organs, including iBAT, iWAT, and gWAT (Guimarães-Camboa et al., 2017). It is conceivable that iBAT and paBAT continue to generate adipocytes during adulthood, but their origins may differ from those observed in the scBAT, tPVAT, and prBAT. These observations elucidate the dynamic nature of brown adipocyte production during adulthood, suggesting distinct mechanisms and potential depot-specific regulations.

Notably, individuals with obesity commonly exhibit reduced BAT activity and a process known as BAT whitening, where brown adipocytes acquire white adipocyte characteristics (Cinti, 2022; Cinti et al., 1997; Jung et al., 1979; Kotzbeck et al., 2018). However, it remains uncertain whether brown adipocytes in these instances retain their original characteristics or if they are replaced by white adipocytes during the whitening process. Using both APC and brown adipocyte lineage tracing tools, it will be intriguing to investigate whether brown adipocytes continue to generate within various depots during obesity and, if so, whether they utilize the same or different cellular sources compared to those observed during adult homeostasis. Based on our current lineage tracing data, we have observed that the contribution of PDGFRβ+ pericytes to brown adipocytes in both PDGFRβ^Rbpj^ and PDGFRβ^RFP^ mice remains relatively stable during the postnatal stage (P10-P30), regardless of the HFHS diet. These findings suggest that the HFHS diet does not significantly impact the involvement of PDGFRβ+ pericytes in the generation of brown adipocytes during this developmental period. While our current study focused on brown adipogenesis under homeostatic conditions in both developmental and adult stages, there is a need to explore how BAT APCs respond to brown adipogenic cues, such as cold challenges and β3 adrenergic receptor agonist CL316,243. For instance, PDGFRα+ stromal cells previously have been shown to contribute to both brown and beige adipogenesis upon cold challenge or CL316,243 treatment (Bukowiecki et al., 1982; Burl et al., 2022; Lee et al., 2015). Additionally, research from Dr. Patrick Seale’s group has identified that brown adipocytes in the PVAT originate from the PDGFRα+ lineage (Angueira et al., 2021). However, there is currently a divergent view regarding whether stromal cells or pericytes serve as the actual APCs. One potential explanation for this divergence could be the co-expression of PDGFRβ and PDGFRα in certain cell populations. A recent study utilizing an inducible dual labeling reporter mouse model demonstrates that cold-induced beige adipogenesis arises partially from PDGFRα+PDGFRβ+ cells (Han et al., 2021). Further studies are needed to clarify the specific contributions of PDGFRα+ and PDGFRβ+ cells to adipogenesis and to discern the differences between them.

Mechanistically, our data show that the deletion of Rbpj in PDGFRβ+ APCs promotes brown adipogenesis during development, both in vivo and in vitro, in a cell-autonomous manner. Multiple studies have shown that Notch signaling plays a role in browning of white adipocytes as well as thermogenic activity in mature brown adipocytes (Bi et al., 2014; Ye et al., 2022) (Huang et al., 2020; Jiang et al., 2017). In addition, a recent publication by Dr. Berry’s group indicated that PDGFRβ signaling in beige APCs negatively regulates cold-induced beige adipogenesis in adulthood, though in a non-cell-autonomous way (Benvie et al., 2023). Yet, despite numerous in vitro and pharmacological studies (Ba et al., 2012; Garcés et al., 1997; Huang et al., 2010; Nueda et al., 2018; Rodríguez-Cano et al., 2020), its function within brown APCs remains uncertain. Our study suggests that brown adipogenesis is enhanced by the inhibition of Notch signaling, which is associated with downregulation of PDGFRβ. However, the exact mechanisms by which Notch regulates PDGFRβ expression, as well as potential impacts on other brown-adipogenic genes like PGC1a and PRDM16, remain unknown. Our results also imply that the presence of PDGF-DD during the brown adipocyte differentiation process significantly suppresses Pparγ expression, possibly leading to inhibition of both adipogenesis and UCP1 expression. This aligns with a recent study that highlighted PDGFRβ’s regulation of Pparγ activity via posttranslational modification (Shao et al., 2021b). Yet, the mechanism by which PDGFRβ regulates Pparγ expression remains unclear. Further mechanistic studies will be required to fully elucidate the molecular and cellular mechanisms by which Notch signaling governs brown adipogenesis.

From a therapeutic perspective, enhanced BAT activity in adults has been shown to have metabolic benefits. However, it’s unclear if promoting brown adipogenesis during development can yield similar impacts. Here using PDGFRβ^Rbpj^ mouse model, our data highlight that promoting postnatal brown adipogenesis effectively mitigates the development of obesity and diabetes induced by a HFHS diet during both developmental and early adult stages. During the postnatal stage (P10-P30), our study uncovers that PDGFRβ^Rbpj^ mice demonstrate enhanced glucose tolerance and reduced body weight gain when subjected to a HFHS diet, in comparison to the control group. Notably, these improvements are linked to increased BAT weights and improved glucose and lipid metabolism within iBAT. We propose that the observed increases in BAT weight, as well as glucose and lipid metabolism, contribute significantly to the favorable metabolic outcomes observed in response to the HFHS diet. Importantly, we still observe the similar metabolic benefits when the HFHS diet was introduced to later P30-P60 adult stage, emphasizing the therapeutic potential of enhancing BAT formation in countering the detrimental effects of an unhealthy diet. Previously, Dr. Gupta’s group has shown that overexpress *Pparγ* in PDGFRβ+ cells during development provide metabolic benefits against diet-induced-obesity in adult stage through manipulation of the gWAT expansion (Zhang et al., 2022). Here we provide another angle suggesting that promoting de novo brown adipogenesis from PDGFRβ+ cells may provide metabolic benefits in adult stage. Given the global pandemic of childhood obesity, our findings underscore the importance of promoting postnatal brown adipogenesis as a potential intervention to address this widespread issue. Of note, the percentage of RFP+ labeled brown adipocytes in the adult stage did not show a significant difference between PDGFRβ^Rbpj^ and PDGFRβ^RFP^ mice. However, the increased tissue weight observed in PDGFRβ^RFP^ mice suggests a potential increase in the total number of RFP+ brown adipocytes. Similarly, analysis of thermogenic gene expression did not reveal any obvious differences between PDGFRβ^Rbpj^ and PDGFRβ^RFP^ mice. Nevertheless, the increase in BAT weight may indicate that the metabolic benefits observed in PDGFRβ^Rbpj^ mice could be attributed to an overall enhancement in the total thermogenic activity of brown adipocytes. It is important to note, however, that due to the presence of PDGFRβ+ cells throughout the body, we cannot exclude the possibility of contributions from other organs or cell types. While our study focuses on the role of brown adipocytes derived from PDGFRβ+ cells, further investigation is needed to elucidate the potential involvement of other organs or cell populations in mediating the observed metabolic effects.

In summary, we have identified pericytes as brown APCs and the Notch-PDGFRβ axis as an inhibitory regulator for brown adipogenesis. Our work sheds light on previously understudied aspects of brown APCs and their regulatory mechanisms, and opens up new avenues for the development of therapies to combat metabolic disorders associated with obesity and diabetes.

### Limitations of the study

One limitation of our study lies in the unidentified differences between PDGFRβ+ and TBX18+ brown APCs during both developmental and adult stages. Future studies, using single-cell RNA sequencing and spatial transcriptomics, may enable us to determine the unique molecular expression profiles underlying these differences.

## METHODS AND MATERIALS

### Mouse models

Mice were housed at 22 ± 2 °C with a humidity of 35 ± 5% and a 14:10 light:dark cycle with a standard rodent chow diet and water unless otherwise indicated. Rosa26^DTA^ (Stock No: 010527) and Rosa26^RFP^ (Stock No. 007914) mice were obtained from the Jackson Laboratory. Ucp1-Cre^ERT2^ mice were generously provided by Dr. Eric N. Olson (University of Texas Southwestern Medical Center). TBX18-Cre^ERT2^ mice were generously provided by Dr. Sylvia Evans (University of California, San Diego). PDGFRβ-Cre^ERT2^ and Rbpj^floxed^ mice were generously provided by Dr. Jan Kitajewski and Dr. Henar Cuvero (University of Illinois at Chicago). All animal experiments were performed according to procedures reviewed and approved by the Institutional Animal Care and Use Committee of the University of Illinois at Chicago.

### Cell culture

The isolated SVF cells were cultured in DMEM/F12 media (Sigma-Aldrich) supplemented with 10% FBS (Sigma-Aldrich) and 1% Penicillin/Streptomycin (Gibco) at 37 °C in a 5% CO_2_ humidified incubator. For beige and brown adipocyte differentiation, ∼90% confluent cells were induced by DMEM/F12 containing 10% FBS, 0.5 mM isobutylmethylxanthine (Sigma-Aldrich), 1 µg/mL insulin (Sigma-Aldrich), 1 µM dexamethasone (Sigma-Aldrich), 60 µM indomethacin (Sigma-Aldrich), 1 nM triiodo-L-thyronine (Sigma-Aldrich), and 1 µM rosiglitazone (Sigma-Aldrich) for 2 days before switched to maintenance medium of DMEM/F12 containing 10% FBS and 1 µg/mL insulin, 1 nM triiodo-L-thyronine, and 1 µM rosiglitazone every 2 days. For iWAT SVF cell differentiation, minimal differentiation protocol was applied, as specified in the context, using minimal differentiation medium of DMEM/F12 containing 10% FBS and 1% Penicillin/Streptomycin and 1ug/ml insulin. Fresh medium was replaced every 2 days until ready for analysis.

### Method details Tamoxifen treatment

To induce Cre translocation and genetic deletion, mice were injected i.p. with 50 mg/kg body weight of Tamoxifen (TAM; Cayman Chemical) dissolved in sunflower oil (Sigma-Aldrich) one time at the designated age. For Cre translocation *in vitro*, cells were treated with 1uM 4-hydroxy-Tamoxifen (4-OHT, Sigma-Alderich)

### Glucose Tolerance Tests

For the glucose tolerance test, mice were fasted for 6 h, starting 9 a.m. till 5 p.m. and were intraperitoneally injected with 25% glucose dissolved in filtered water (1.5 g/kg). Glucose levels were measured from tail blood at 0, 15, 30, 60, 90 and 120 min after glucose injection by using a glucose meter (Contour Next).

### Isolation of stromal vascular fraction (SVF)

Isolation of SVF cells from iWAT, iBAT, gWAT, scBAT and PVAT was performed as previously described (Park et al., 2021). Briefly, adipose tissue was excised from mice of designated age and minced by scissors and then digested in isolation buffer (100 mM HEPES, 0.12 M NaCl, 50 mM KCl, 5 mM D-glucose, 1.5% BSA, 1 mM CaCl_2_, pH 7.3) containing 1 mg/ml type I collagenase (Worthington) at 37 °C for 45 min. Digested tissue was filtered through 70 µm mesh to remove large pieces, and the flow-through was then centrifuged at 800 × g for 10 min. The pellets were resuspended and seeded. After 1 hour of plating, red blood cell lysis buffer (155 mM NH_4_Cl, 10 mM KHCO_3_, 0.1 mM EDTA) was replaced in the culture plates for 5 mins before replaced back to culture medium.

### Western blot analysis

RIPA buffer was used to isolate protein from cells and centrifuged at 14,000 x g for 15 mins at 4 C. The sample concentration was determined using BCA assay kit (Thermo Fisher Scientific). 10 ug of total protein of each sample was loaded in SDS-PAGE gel for separation and transferred to PVDF membranes. The membranes were blocked with 5% non-fat milk at room temperature followed by primary antibody dissolved in 5% non-fat milk at 4C overnight. After 3 x 10 mins washes with TBST, secondary antibodies in 5% non-fat milk were used to incubate the membranes for another 1 hr at room temperature. The signals were revealed using Western ECL substrate kit with Bio-Rad imager. Antibody information are available in Key Resources Table.

### Histological staining

Hematoxylin and eosin (H&E) staining were performed using standard methods as described in previous publications (Shin et al., 2020). Briefly, tissues were fixed in 10% formalin overnight, dehydrated, embedded in paraffin, and sectioned with a microtome at 4-8 μm thicknesses. For immunofluorescence staining, paraffin sections were incubated with permeabilization buffer (0.3% Triton X-100 in PBS) for 30 min at room temperature, with primary antibody at 4°C overnight, and with secondary antibody for 2 h at room temperature, all in blocking buffer (5% donkey serum in 1X PBS). Images were taken using a confocal laser microscope (Carl Zeiss Ltd). Adipocyte sizes were quantified by using Image J.

### Oil red O staining

Cells were washed with PBS and fixed in 4% PFA at room temperature for 30 mins. After washing with PBS three times, cells were stained with 60% filtered Oil Red O working solution (vol/vol in distilled water) at room temperature for 30 minutes. Cells then were washed with ddH_2_O three times, 10 minutes each before imaging. Lipid accumulation was quantified by eluting Oil Red O trapped in cells with 100% isopropanol and then analyzed at 500 nm in a plate reader.

### RNA isolation and quantitative real-time PCR (qPCR) analysis

Total RNA was isolated from cells or tissues using Tripure Isolation Reagent (Sigma-Aldrich) and Bullet Blender Homogenizer (Next Advance) following manufacturer’s protocol. 1 ug of isolated RNA was reverse transcribed to cDNA using cDNA reverse-transcription kit (Applied Biosystems). 2X Syber Green qPCR master mix (Abclonal) was used for qPCR analysis following manufacturer’s instructions and analyzed on a ViiA7 system (Applied Biosystems). Data were analyzed using comparative CT methods. The relative expression of mRNAs was normalized to actin. Primer sequences are available in Table S1.

### Immunofluorescent staining

Sectioned slides were incubated in 55 C° over night to remove access paraffin. Slides were chilled before going through 3 x 3 mins xylene, 3 x 1 min 100% ethanol, 2 x 1min 95% ethanol and 1 x 3 mins ddH_2_O for rehydration. Rehydrated slides went through antigen retrieval buffer (1mM Tris-HCL, pH=8) in a microwave till boiling point in coplin jars. Slides were then cooled to room temperature before washed with PBS and blocked with blocking buffer (10% Donkey Serum in PBS) for 1 hr followed by primary antibody incubation over night at 4°C. Sections were then washed with PBS 3 x 10 mins before secondary antibody incubation for 1 hour at room temperature followed by another PBS 3 x 10 mins wash. Sections were then mounted with coverslips for imaging. Coloc2 plugin in ImageJ was used for quantification of colocalization of PLIN and RFP in pixels. And Mander’s coefficient was used for RFP+PLIN+/PLIN+ pixels.

### Statistical analysis

Studies were performed on two or three independent cohorts and were performed on 6-8 mice per group unless specified. Data are presented as mean ± SEM. Two-tailed unpaired Student’s t test (for comparison of two groups) or one-way ANOVA followed by Tukey’s test (for comparison of three or more groups) were conducted using GraphPad Prism software. Asterisks indicate statistically significant differences between groups. *, P ≤ 0.05; **, P ≤ 0.01; and ***, P ≤ 0.001, ****, P ≤ 0.0001.

## Supporting information

KEY RESOURCES TABLE

## ACKNOWLEDGMENTS

We thank Dr. Lijun Rong for supporting this work and giving constructive suggestions. We thank Dr. Sylvia Evans and Dr. Henar Cuervo for kindly providing the TBX18-Cre^ERT2^ and PDGFRβ-Cre^ERT2^ mouse strains. We thank Cynthia Rose Adams and Jeanette Purcell for assistance with mouse husbandry, Stefan J. Green and the Research Resources Center for RT-PCR analysis, Dr. Brian Layden and Metabolic Phenotyping Core for analytical and phenotypical mouse measurements and members of the Jiang laboratory for helpful comments on the manuscript. This work was supported by grants to Y.J. from the National Institute of Diabetes and Digestive and Kidney Disease grant (NIDDK) K01 DK11177, R03 DK127149, R01 DK132398-02 and Pilot & Feasibility Diabetes Research & Training Center (DRTC) Award (P30DK020595).

## AUTHOR CONTRIBUTIONS

Y.J. and Z.S. conceived and designed the experiments. Z.S. and S.X. conducted most of the experiments. Z.S., S.X. and Y.J. interpreted the experiments. Z.S. and Y.J. wrote and revised the manuscript, and Z.S., S.X., R.H., Z.W., J.P., Y.Q., J.W., P.B., N.V., Q.S., Z.S., B.L., and Y.J. all authors reviewed it.

## DATA AVAILABILITY STATEMENT

Not applicable

---

**Supplementary Figure 1.**
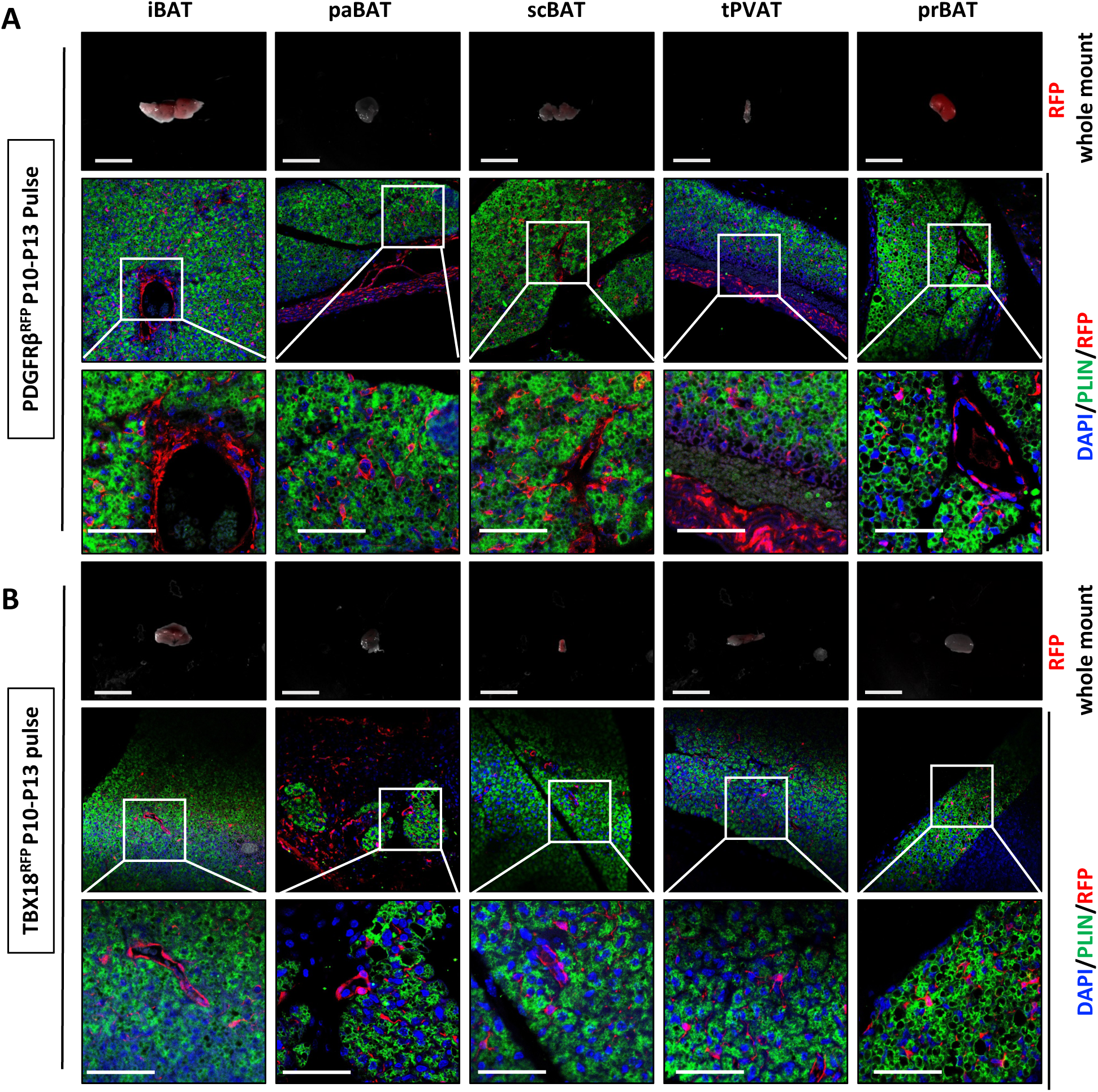
PDGFRβ^RFP^ and TBX18^RFP^ faithfully label pericytes at pulse. **(A)** Upper panel: Whole mount RFP images of PDGFRβ^RFP^ BAT depots at P13. Scale bar, 1mm. Middle and lower panel: IF staining of DAPI (blue), PLIN (green) and RFP (red). Lower Panel: Zoomed-in images of middle panel. Scale Bar, 50 µm. **(B)** Upper panel: Whole mount RFP images of TBX18^RFP^ BAT depots at P13. Scale bar, 1mm. Middle and lower panel: IF staining of DAPI (blue), PLIN (green) and RFP (red). Lower Panel: Zoomed-in images of middle panel. Scale Bar, 50 µm.

**Supplementary Figure 2.**
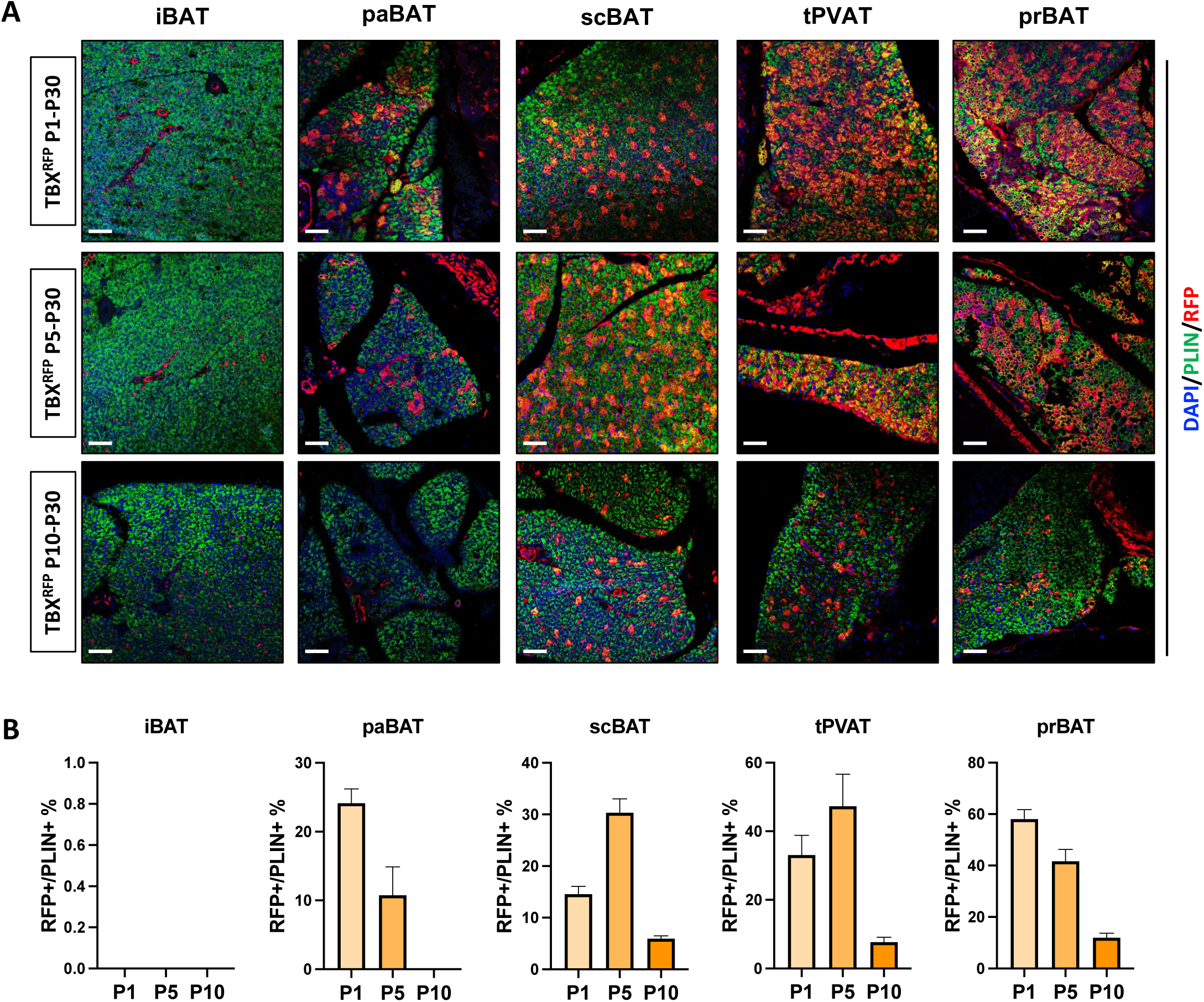
Lineage tracing of TBX18^RFP^ mice in early postnatal stage. **(A)** Representative images of IF staining of DAPI (blue), PLIN (green) and RFP (red). Scale bar, 50 µm. **(B)** Quantification of colocalization of PLIN and RFP of IF staining by pixels (n=3-4 images). Data are presented as means ± SEM.

**Supplementary Figure 3.**
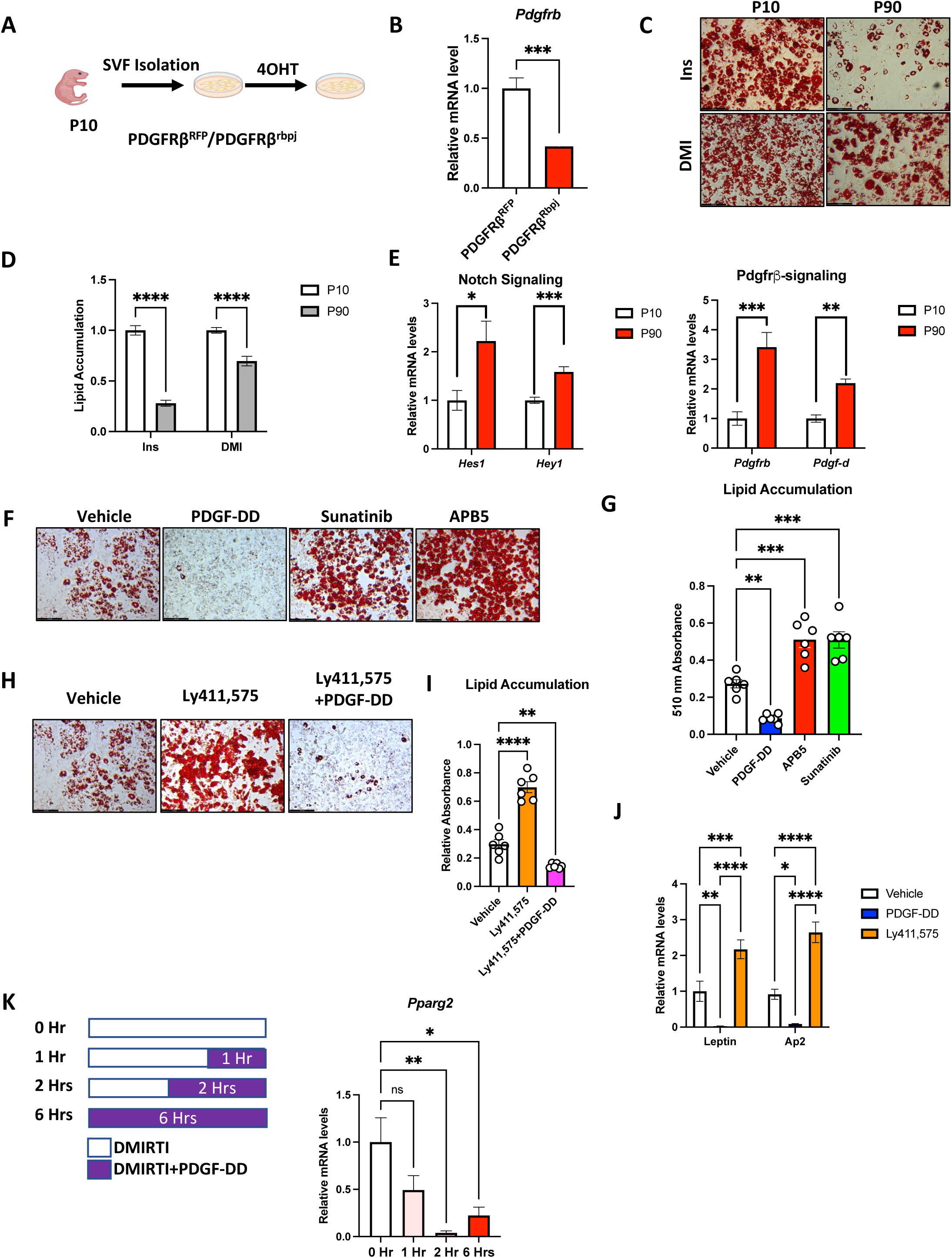
Notch inhibition promoted brown adipogenesis is reversed by Pdgfrβ downregulation. **(A)** Experimental Design. SVF, isolated from PDGFRβ^RFP^ and PDGFRβ^Rbpj^ mice, are treated with 4OHT to induce Rbpj knockout. **(B)** qRT-PCR analysis of Pdgfrb from PDGFRβ^RFP^ and PDGFRβ^Rbpj^ SVF. (n=6 per group). ***p<0.001. Data are presented as means ± SEM. **(C)** Oil-Red O staining of P10 and P90 SVF differentiated with either insulin alone or full differentiation cocktail. **(D)** Oil-Red O accumulation quantification of (C) (n=5 per group). ****p<0.0001. Data are presented as means ± SEM. **(E)** qRT-PCR analysis of Notch and Pdgfrβ signaling genes of P10 and P90 SVF (n=6 per group). *p<0.05, **p<0.01, ***p<0.01. Data are presented as means ± SEM. **(F)** Oil-Red O staining of P10 SVF with drug pretreatment and quantification. **(G)** Oil-Red O staining of P10 SVF with drug pretreatment and quantification SVF (n=6 per group). **p<0.01, ***p<0.01. Data are presented as means ± SEM. **(H)** Oil-Red O staining of P10 SVF with drug pretreatment and quantification. **(I)** Oil-Red O staining of P10 SVF with drug pretreatment and quantification SVF (n=6 per group). **p<0.01, ***p<0.01. Data are presented as means ± SEM. **(J)** qRT-PCR analysis of adipogenic genes (n=5 per group). *p<0.05, **p<0.01, ***p<0.01, ****p<0.01. Data are presented as means ± SEM. **(K)** Left panel: experimental design. Right panel: qRT-PCR analysis of *Pparg2* (n=4 per group). *p<0.05, **p<0.01. Data are presented as means ± SEM. DMIRTI is the beige/brown differentiation cocktail, D-dexamethasone, M-IBMX, I-insulin, R-Rosiglitazone, T-triiodo-L-thyronine, I-Indomethacin.

**Supplementary Figure 4.**
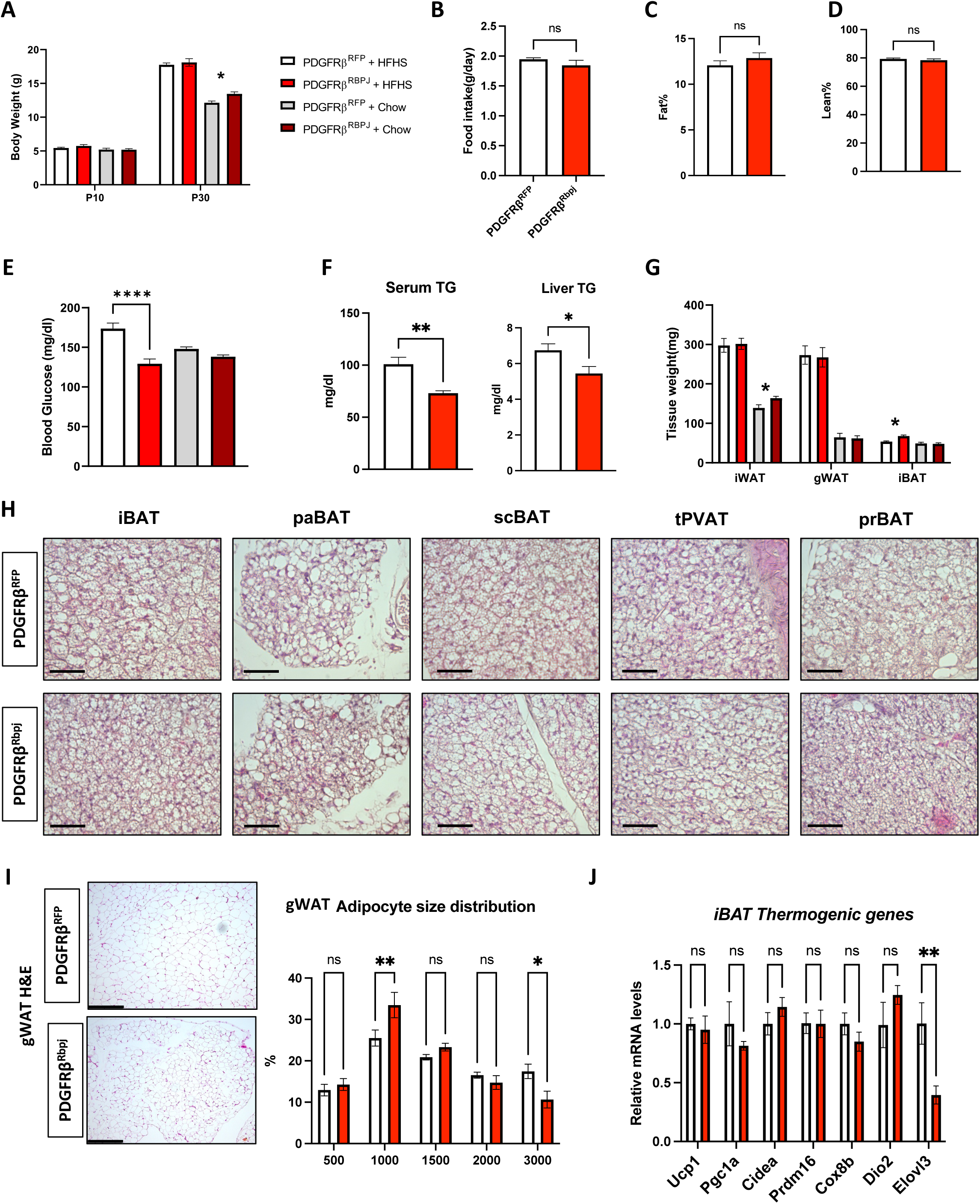
Notch inhibition in PDGFRβ+ cells prevents diet-induced metabolic stress in postnatal stage. **(A)** Body weight of PDGFRβ^RFP^ and PDGFRβ^Rbpj^ mice fed on chow or HFHS diet (n>10 per group). *p< 0.05. Data are presented as means ± SEM. Food intake. Data are presented as means ± SEM. **(B)** Fat content. Data are presented as means ± SEM. **(C)** Lean content. Data are presented as means ± SEM. **(D)** Fed blood glucose. ****p< 0.0001. Data are presented as means ± SEM. **(E)** Serum and liver triglyceride level. *p< 0.05, **p< 0.01. Data are presented as means ± SEM. **(F)** Adipose tissue weights. *p< 0.05. Data are presented as means ± SEM. **(G)** H&E staining of BAT depots of HFHS fed PDGFRβ^RFP^ and PDGFRβ^Rbpj^ mice. Scale bar, 50 µm. **(H)** Representative H&E images of gWAT. Adipocyte size distribution analysis. *p< 0.05. Data are presented as means ± SEM. **(I)** qRT-PCR analysis of thermogenic genes of iBAT. **p< 0.01. Data are presented as means ± SEM.

**Supplementary Figure 5.**
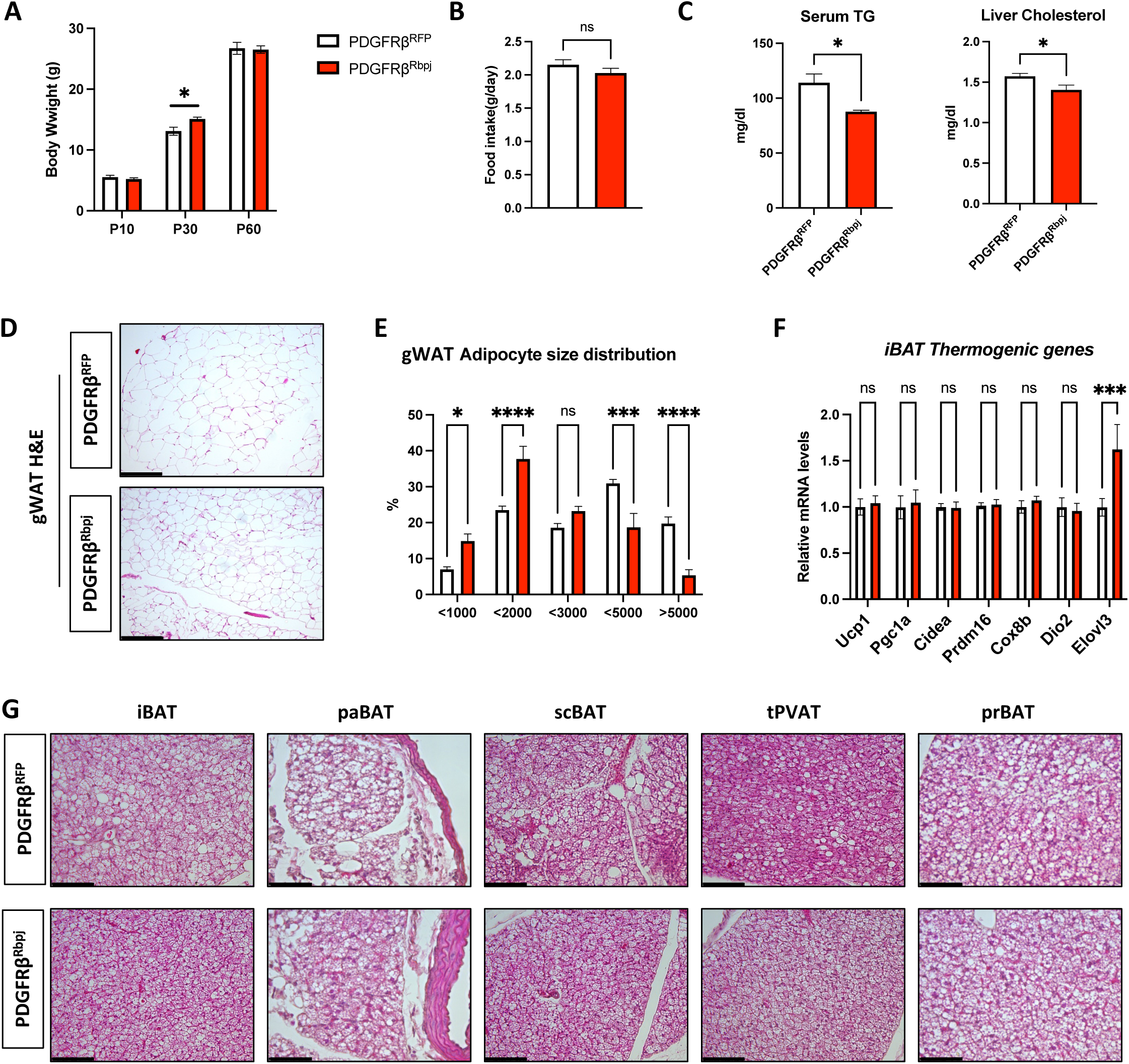
Notch inhibition in PDGFRβ+ cells prevents diet-induced metabolic stress in early adult stage. **(A)** Body weight of PDGFRβ^RFP^ and PDGFRβ^Rbpj^ mice fed on HFHFS deit. *p< 0.05. Data are presented as means ± SEM. **(B)** Food intake. Data are presented as means ± SEM. **(C)** Serum triglyceride and liver cholesterol level. *p< 0.05. Data are presented as means ± SEM. **(D)** Serum TG and liver cholesterol level. **(E)** Representative H&E images of gWAT. Scale bar, 50 µm. **(F)** Adipocyte size distribution analysis. *p< 0.05, ***p< 0.001, ****p< 0.0001. Data are presented as means ± SEM. **(G)** qRT-PCR analysis of thermogenic genes of iBAT. **p< 0.01. Data are presented as means ± SEM. **(H)** H&E staining of BAT depots of HFHS fed PDGFRβ^RFP^ and PDGFRβ^Rbpj^ mice. Scale bar, 50 µm.

## REFERENCES

1. Adachi, Y., Ueda, K., Nomura, S., Ito, K., Katoh, M., Katagiri, M., Yamada, S., Hashimoto, M., Zhai, B., Numata, G., et al. (2022). Beiging of perivascular adipose tissue regulates its inflammation and vascular remodeling. Nat Commun 13, 5117.

2. Afshin, A., Forouzanfar, M.H., Reitsma, M.B., Sur, P., Estep, K., Lee, A., Marczak, L., Mokdad, A.H., Moradi-Lakeh, M., Naghavi, M., et al. (2017). Health Effects of Overweight and Obesity in 195 Countries over 25 Years. N Engl J Med 377, 13–27.

3. Angueira, A.R., Sakers, A.P., Holman, C.D., Cheng, L., Arbocco, M.N., Shamsi, F., Lynes, M.D., Shrestha, R., Okada, C., Batmanov, K., et al. (2021). Defining the lineage of thermogenic perivascular adipose tissue. Nat Metab 3, 469–484.

4. Ba, K., Yang, X., Wu, L., Wei, X., Fu, N., Fu, Y., Cai, X., Yao, Y., Ge, Y., and Lin, Y. (2012). Jagged-1-mediated activation of notch signalling induces adipogenesis of adipose-derived stem cells. Cell Prolif 45, 538–544.

5. Barandier, C., Montani, J.P., and Yang, Z. (2005). Mature adipocytes and perivascular adipose tissue stimulate vascular smooth muscle cell proliferation: effects of aging and obesity. Am J Physiol Heart Circ Physiol 289, H1807–1813.

6. Becher, T., Palanisamy, S., Kramer, D.J., Eljalby, M., Marx, S.J., Wibmer, A.G., Butler, S.D., Jiang, C.S., Vaughan, R., Schoder, H., et al. (2021). Brown adipose tissue is associated with cardiometabolic health. Nat Med 27, 58–65.

7. Benvie, A.M., Lee, D., Steiner, B.M., Xue, S., Jiang, Y., and Berry, D.C. (2023). Age-dependent Pdgfrbeta signaling drives adipocyte progenitor dysfunction to alter the beige adipogenic niche in male mice. Nat Commun 14, 1806.

8. Berry, D.C., Jiang, Y., and Graff, J.M. (2016). Mouse strains to study cold-inducible beige progenitors and beige adipocyte formation and function. Nat Commun 7, 10184.

9. Betz, M.J., and Enerback, S. (2011). Therapeutic prospects of metabolically active brown adipose tissue in humans. Front Endocrinol (Lausanne) 2, 86.

10. Bi, P., Shan, T., Liu, W., Yue, F., Yang, X., Liang, X.R., Wang, J., Li, J., Carlesso, N., Liu, X., et al. (2014). Inhibition of Notch signaling promotes browning of white adipose tissue and ameliorates obesity. Nat Med 20, 911–918.

11. Bi, P., Yue, F., Karki, A., Castro, B., Wirbisky, S.E., Wang, C., Durkes, A., Elzey, B.D., Andrisani, O.M., Bidwell, C.A., et al. (2016). Notch activation drives adipocyte dedifferentiation and tumorigenic transformation in mice. J Exp Med 213, 2019–2037.

12. Boucher, J.M., Ryzhova, L., Harrington, A., Davis-Knowlton, J., Turner, J.E., Cooper, E., Maridas, D., Ryzhov, S., Rosen, C.J., Vary, C.P.H., et al. (2020). Pathological Conversion of Mouse Perivascular Adipose Tissue by Notch Activation. Arterioscler Thromb Vasc Biol 40, 2227–2243.

13. Bukowiecki, L., Collet, A.J., Follea, N., Guay, G., and Jahjah, L. (1982). Brown adipose tissue hyperplasia: a fundamental mechanism of adaptation to cold and hyperphagia. Am J Physiol 242, E353–359.

14. Burl, R.B., Rondini, E.A., Wei, H., Pique-Regi, R., and Granneman, J.G. (2022). Deconstructing cold-induced brown adipocyte neogenesis in mice. Elife 11.

15. Cai, Y.Y., Zhang, H.B., Fan, C.X., Zeng, Y.M., Zou, S.Z., Wu, C.Y., Wang, L., Fang, S., Li, P., Xue, Y.M., et al. (2019). Renoprotective effects of brown adipose tissue activation in diabetic mice. J Diabetes 11, 958–970.

16. Castel, D., Mourikis, P., Bartels, S.J., Brinkman, A.B., Tajbakhsh, S., and Stunnenberg, H.G. (2013). Dynamic binding of RBPJ is determined by Notch signaling status. Genes Dev 27, 1059–1071.

17. Cawthorn, W.P., Scheller, E.L., and MacDougald, O.A. (2012). Adipose tissue stem cells: the great WAT hope. Trends Endocrinol Metab 23, 270–277.

18. Cheng, Y.W., Zhang, Z.B., Lan, B.D., Lin, J.R., Chen, X.H., Kong, L.R., Xu, L., Ruan, C.C., and Gao, P.J. (2021). PDGF-D activation by macrophage-derived uPA promotes AngII-induced cardiac remodeling in obese mice. J Exp Med 218.

19. Chi, J., Lin, Z., Barr, W., Crane, A., Zhu, X.G., and Cohen, P. (2021). Early postnatal interactions between beige adipocytes and sympathetic neurites regulate innervation of subcutaneous fat. Elife 10.

20. Chondronikola, M., Volpi, E., Børsheim, E., Porter, C., Saraf, M.K., Annamalai, P., Yfanti, C., Chao, T., Wong, D., Shinoda, K., et al. (2016). Brown Adipose Tissue Activation Is Linked to Distinct Systemic Effects on Lipid Metabolism in Humans. Cell Metab 23, 1200–1206.

21. Cinti, S. (2022). The endocrine adipose organ. Rev Endocr Metab Disord 23, 1–4.

22. Cinti, S., Frederich, R.C., Zingaretti, M.C., De Matteis, R., Flier, J.S., and Lowell, B.B. (1997). Immunohistochemical localization of leptin and uncoupling protein in white and brown adipose tissue. Endocrinology 138, 797–804.

23. Cittadini, A., Mantzoros, C.S., Hampton, T.G., Travers, K.E., Katz, S.E., Morgan, J.P., Flier, J.S., and Douglas, P.S. (1999). Cardiovascular abnormalities in transgenic mice with reduced brown fat: an animal model of human obesity. Circulation 100, 2177–2183.

24. Entringer, S., Rasmussen, J., Cooper, D.M., Ikenoue, S., Waffarn, F., Wadhwa, P.D., and Buss, C. (2017). Association between supraclavicular brown adipose tissue composition at birth and adiposity gain from birth to 6 months of age. Pediatr Res 82, 1017–1021.

25. Fu, M., Xu, L., Chen, X., Han, W., Ruan, C., Li, J., Cai, C., Ye, M., and Gao, P. (2019). Neural Crest Cells Differentiate Into Brown Adipocytes and Contribute to Periaortic Arch Adipose Tissue Formation. Arterioscler Thromb Vasc Biol 39, 1629–1644.

26. Garcés, C., Ruiz-Hidalgo, M.J., Font de Mora, J., Park, C., Miele, L., Goldstein, J., Bonvini, E., Porrás, A., and Laborda, J. (1997). Notch-1 controls the expression of fatty acid-activated transcription factors and is required for adipogenesis. J Biol Chem 272, 29729–29734.

27. Guimarães-Camboa, N., Cattaneo, P., Sun, Y., Moore-Morris, T., Gu, Y., Dalton, N.D., Rockenstein, E., Masliah, E., Peterson, K.L., Stallcup, W.B., et al. (2017). Pericytes of Multiple Organs Do Not Behave as Mesenchymal Stem Cells In Vivo. Cell Stem Cell 20, 345–359.e345.

28. Gupta, R.K., Mepani, R.J., Kleiner, S., Lo, J.C., Khandekar, M.J., Cohen, P., Frontini, A., Bhowmick, D.C., Ye, L., Cinti, S., et al. (2012). Zfp423 expression identifies committed preadipocytes and localizes to adipose endothelial and perivascular cells. Cell Metab 15, 230–239.

29. Han, X., Zhang, Z., He, L., Zhu, H., Li, Y., Pu, W., Han, M., Zhao, H., Liu, K., Huang, X., et al. (2021). A suite of new Dre recombinase drivers markedly expands the ability to perform intersectional genetic targeting. Cell Stem Cell 28, 1160–1176.e1167.

30. Huang, C.L., Huang, Y.N., Yao, L., Li, J.P., Zhang, Z.H., Huang, Z.Q., Chen, S.X., Zhang, Y.L., Wang, J.F., Chen, Y.X., et al. (2023). Thoracic perivascular adipose tissue inhibits VSMC apoptosis and aortic aneurysm formation in mice via the secretome of browning adipocytes. Acta Pharmacol Sin 44, 345–355.

31. Huang, D., Narayanan, N., Cano-Vega, M.A., Jia, Z., Ajuwon, K.M., Kuang, S., and Deng, M. (2020). Nanoparticle-Mediated Inhibition of Notch Signaling Promotes Mitochondrial Biogenesis and Reduces Subcutaneous Adipose Tissue Expansion in Pigs. iScience 23, 101167.

32. Huang, Y., Yang, X., Wu, Y., Jing, W., Cai, X., Tang, W., Liu, L., Liu, Y., Grottkau, B.E., and Lin, Y. (2010). gamma-secretase inhibitor induces adipogenesis of adipose-derived stem cells by regulation of Notch and PPAR-gamma. Cell Prolif 43, 147–156.

33. Jiang, C., Cano-Vega, M.A., Yue, F., Kuang, L., Narayanan, N., Uzunalli, G., Merkel, M.P., Kuang, S., and Deng, M. (2017). Dibenzazepine-Loaded Nanoparticles Induce Local Browning of White Adipose Tissue to Counteract Obesity. Mol Ther 25, 1718–1729.

34. Jiang, Y., Berry, D.C., Tang, W., and Graff, J.M. (2014). Independent stem cell lineages regulate adipose organogenesis and adipose homeostasis. Cell Rep 9, 1007–1022.

35. Jin, S., Hansson, E.M., Tikka, S., Lanner, F., Sahlgren, C., Farnebo, F., Baumann, M., Kalimo, H., and Lendahl, U. (2008). Notch signaling regulates platelet-derived growth factor receptor-beta expression in vascular smooth muscle cells. Circ Res 102, 1483–1491.

36. Jung, R.T., Shetty, P.S., James, W.P., Barrand, M.A., and Callingham, B.A. (1979). Reduced thermogenesis in obesity. Nature 279, 322–323.

37. Kangsamaksin, T., Tattersall, I.W., and Kitajewski, J. (2014). Notch functions in developmental and tumour angiogenesis by diverse mechanisms. Biochem Soc Trans 42, 1563–1568.

38. Kotzbeck, P., Giordano, A., Mondini, E., Murano, I., Severi, I., Venema, W., Cecchini, M.P., Kershaw, E.E., Barbatelli, G., Haemmerle, G., et al. (2018). Brown adipose tissue whitening leads to brown adipocyte death and adipose tissue inflammation. J Lipid Res 59, 784–794.

39. Lee, Y.H., Petkova, A.P., Konkar, A.A., and Granneman, J.G. (2015). Cellular origins of cold-induced brown adipocytes in adult mice. FASEB J 29, 286–299.

40. Lepper, C., and Fan, C.M. (2010). Inducible lineage tracing of Pax7-descendant cells reveals embryonic origin of adult satellite cells. Genesis 48, 424–436.

41. Li, X., Ballantyne, L.L., Yu, Y., and Funk, C.D. (2019). Perivascular adipose tissue-derived extracellular vesicle miR-221-3p mediates vascular remodeling. FASEB J 33, 12704–12722.

42. Liu, X., Wang, S., You, Y., Meng, M., Zheng, Z., Dong, M., Lin, J., Zhao, Q., Zhang, C., Yuan, X., et al. (2015). Brown Adipose Tissue Transplantation Reverses Obesity in Ob/Ob Mice. Endocrinology 156, 2461–2469.

43. Mayeuf-Louchart, A., Lancel, S., Sebti, Y., Pourcet, B., Loyens, A., Delhaye, S., Duhem, C., Beauchamp, J., Ferri, L., Thorel, Q., et al. (2019). Glycogen Dynamics Drives Lipid Droplet Biogenesis during Brown Adipocyte Differentiation. Cell Rep 29, 1410–1418.e1416.

44. Min, S.Y., Kady, J., Nam, M., Rojas-Rodriguez, R., Berkenwald, A., Kim, J.H., Noh, H.L., Kim, J.K., Cooper, M.P., Fitzgibbons, T., et al. (2016). Human ‘brite/beige’ adipocytes develop from capillary networks, and their implantation improves metabolic homeostasis in mice. Nat Med 22, 312–318.

45. Mo, Q., Salley, J., Roshan, T., Baer, L.A., May, F.J., Jaehnig, E.J., Lehnig, A.C., Guo, X., Tong, Q., Nuotio-Antar, A.M., et al. (2017). Identification and characterization of a supraclavicular brown adipose tissue in mice. JCI Insight 2.

46. Nadeem, T., Bogue, W., Bigit, B., and Cuervo, H. (2020). Deficiency of Notch signaling in pericytes results in arteriovenous malformations. JCI Insight 5.

47. Nueda, M.L., González-Gómez, M.J., Rodríguez-Cano, M.M., Monsalve, E.M., Díaz-Guerra, M.J.M., Sánchez-Solana, B., Laborda, J., and Baladrón, V. (2018). DLK proteins modulate NOTCH signaling to influence a brown or white 3T3-L1 adipocyte fate. Sci Rep 8, 16923.

48. Park, J., Shin, S., Liu, L., Jahan, I., Ong, S.G., Xu, P., Berry, D.C., and Jiang, Y. (2021). Progenitor-like characteristics in a subgroup of UCP1+ cells within white adipose tissue. Dev Cell 56, 985–999 e984.

49. Rodríguez-Cano, M.M., González-Gómez, M.J., Sánchez-Solana, B., Monsalve, E.M., Díaz-Guerra, M.M., Laborda, J., Nueda, M.L., and Baladrón, V. (2020). NOTCH Receptors and DLK Proteins Enhance Brown Adipogenesis in Mesenchymal C3H10T1/2 Cells. Cells 9.

50. Seki, T., Yang, Y., Sun, X., Lim, S., Xie, S., Guo, Z., Xiong, W., Kuroda, M., Sakaue, H., Hosaka, K., et al. (2022). Brown-fat-mediated tumour suppression by cold-altered global metabolism. Nature 608, 421–428.

51. Shamsi, F., Wang, C.H., and Tseng, Y.H. (2021). The evolving view of thermogenic adipocytes – ontogeny, niche and function. Nat Rev Endocrinol 17, 726–744.

52. Shao, M., Hepler, C., Zhang, Q., Shan, B., Vishvanath, L., Henry, G.H., Zhao, S., An, Y.A., Wu, Y., Strand, D.W., et al. (2021a). Pathologic HIF1alpha signaling drives adipose progenitor dysfunction in obesity. Cell Stem Cell 28, 685–701 e687.

53. Shao, M., Hepler, C., Zhang, Q., Shan, B., Vishvanath, L., Henry, G.H., Zhao, S., An, Y.A., Wu, Y., Strand, D.W., et al. (2021b). Pathologic HIF1α signaling drives adipose progenitor dysfunction in obesity. Cell Stem Cell 28, 685–701.e687.

54. Shin, S., Pang, Y., Park, J., Liu, L., Lukas, B.E., Kim, S.H., Kim, K.W., Xu, P., Berry, D.C., and Jiang, Y. (2020). Dynamic control of adipose tissue development and adult tissue homeostasis by platelet-derived growth factor receptor alpha. Elife 9.

55. Song, A., Dai, W., Jang, M.J., Medrano, L., Li, Z., Zhao, H., Shao, M., Tan, J., Li, A., Ning, T., et al. (2020). Low-and high-thermogenic brown adipocyte subpopulations coexist in murine adipose tissue. J Clin Invest 130, 247–257.

56. Stefan, N., Häring, H.U., and Cusi, K. (2019). Non-alcoholic fatty liver disease: causes, diagnosis, cardiometabolic consequences, and treatment strategies. Lancet Diabetes Endocrinol 7, 313–324.

57. Takaoka, M., Nagata, D., Kihara, S., Shimomura, I., Kimura, Y., Tabata, Y., Saito, Y., Nagai, R., and Sata, M. (2009). Periadventitial adipose tissue plays a critical role in vascular remodeling. Circ Res 105, 906–911.

58. Tam, C.S., Lecoultre, V., and Ravussin, E. (2012). Brown adipose tissue: mechanisms and potential therapeutic targets. Circulation 125, 2782–2791.

59. Tang, W., Zeve, D., Suh, J.M., Bosnakovski, D., Kyba, M., Hammer, R.E., Tallquist, M.D., and Graff, J.M. (2008). White fat progenitor cells reside in the adipose vasculature. Science 322, 583–586.

60. Thoonen, R., Ernande, L., Cheng, J., Nagasaka, Y., Yao, V., Miranda-Bezerra, A., Chen, C., Chao, W., Panagia, M., Sosnovik, D.E., et al. (2015). Functional brown adipose tissue limits cardiomyocyte injury and adverse remodeling in catecholamine-induced cardiomyopathy. J Mol Cell Cardiol 84, 202–211.

61. Vishvanath, L., MacPherson, K.A., Hepler, C., Wang, Q.A., Shao, M., Spurgin, S.B., Wang, M.Y., Kusminski, C.M., Morley, T.S., and Gupta, R.K. (2015). Pdgfrbeta Mural Preadipocytes Contribute to Adipocyte Hyperplasia Induced by High-Fat-Diet Feeding and Prolonged Cold Exposure in Adult Mice. Cell Metab.

62. Vishvanath, L., MacPherson, K.A., Hepler, C., Wang, Q.A., Shao, M., Spurgin, S.B., Wang, M.Y., Kusminski, C.M., Morley, T.S., and Gupta, R.K. (2016). Pdgfrbeta+ Mural Preadipocytes Contribute to Adipocyte Hyperplasia Induced by High-Fat-Diet Feeding and Prolonged Cold Exposure in Adult Mice. Cell Metab 23, 350-359.

63. Wang, Y., Gao, M., Zhu, F., Li, X., Yang, Y., Yan, Q., Jia, L., Xie, L., and Chen, Z. (2020). METTL3 is essential for postnatal development of brown adipose tissue and energy expenditure in mice. Nat Commun 11, 1648.

64. Ye, M.S., Luo, L., Guo, Q., Su, T., Cheng, P., and Huang, Y. (2022). KCTD10 regulates brown adipose tissue thermogenesis and metabolic function via Notch signaling. J Endocrinol 252, 155–166.

65. Zhang, F., Hao, G., Shao, M., Nham, K., An, Y., Wang, Q., Zhu, Y., Kusminski, C.M., Hassan, G., Gupta, R.K., et al. (2018a). An Adipose Tissue Atlas: An Image-Guided Identification of Human-like BAT and Beige Depots in Rodents. Cell Metab 27, 252–262 e253.

66. Zhang, Q., Shan, B., Guo, L., Shao, M., Vishvanath, L., Elmquist, G., Xu, L., and Gupta, R.K. (2022). Distinct functional properties of murine perinatal and adult adipose progenitor subpopulations. Nat Metab 4, 1055–1070.

67. Zhang, Y., Cai, Y., Zhang, H., Zhang, J., Zeng, Y., Fan, C., Zou, S., Wu, C., Fang, S., Li, P., et al. (2021). Brown adipose tissue transplantation ameliorates diabetic nephropathy through the miR-30b pathway by targeting Runx1. Metabolism 125, 154916.

68. Zhang, Z.B., Ruan, C.C., Lin, J.R., Xu, L., Chen, X.H., Du, Y.N., Fu, M.X., Kong, L.R., Zhu, D.L., and Gao, P.J. (2018b). Perivascular Adipose Tissue-Derived PDGF-D Contributes to Aortic Aneurysm Formation During Obesity. Diabetes 67, 1549–1560.

