## Supplementary material for "The Notch-Pdgfrβ axis suppresses brown adipocyte progenitor differentiation in early postnatal mice": KEY RESOURCES TABLE

**STAR METHODS**
**KEY RESOURCES TABLE**

| REAGENT or RESOURCE | SOURCE | IDENTIFIER |
|---|---|---|
| Antibodies | | |
| Rabbit anti-UCP1 | Thermo Scientific | Cat# PA1-24894 |
| Rabbit anti-Rbpj | Cell Signaling Technology | Cat# 5442 |
| Rabbit anti-RFP | Novus Biologicals | Cat# NBP1-69962 |
| Rabbit anti-ACTB | Cell Signaling Technology | Cat# 4970 |
| Goat anti-Perilipin | Abcam | Cat#: ab61682 |
| HRP goat anti-rabbit | Cell Signaling Technology | Cat#: 96714S |
| Cd140b Monoclonal Antibody (APB5) | eBioscience | Cat#: 14-1402-82 |
| Cy5 donkey anti-goat | Jackson ImmunoResearch | Cat#: 705-175-147 |
| Cy5 donkey anti-rabbit | Jackson ImmunoResearch | Cat#: 711-175-152 |
| Chemicals, Peptides, and Recombinant Proteins | | |
| Collagenase, Type 1 | Worthington Biochemical | Cat#: LS004197 |
| Dexamethasone | Sigma-Aldrich | Cat#: D4902 |
| Fetal Bovine Serum | Corning | Cat#: 35-015-CV |
| Glucose | Sigma-Aldrich | Cat#: G7021-100G |
| Indomethacin | Sigma-Aldrich | Cat#: 53-86-1 |
| Isobutylmethylxanthine (IBMX) | Sigma-Aldrich | Cat#: I5879 |
| triiodo-L-thyronine (T3) | Sigma-Aldrich | Cat#: T2877 |
| Insulin | Sigma-Aldrich | Cat#: I6634 |
| 4-hydroxy-Tamoxifen | Sigma-Aldrich | Cat#: H7904 |
| DAPI | Sigma-Aldrich | Cat#: D9564 |
| Tamoxifen | Sigma-Aldrich | Cat#: T5648 |
| Protease Inhibitor Cocktail | MedChem Express | Cat#: HY-K0011 |
| Bovine Serum Albumin | Sigma-Aldrich | Cat#: A9418 |
| Rosiglitazone | Sigma-Aldrich | Cat#: R-2408 |
| Recombinant Mouse PDGF-D Protein, CF | R&D Systems | Cat#: 9738-SB-050 |
| Ly411575 | VWR Selleck Chemicals | Cat#: S2714 |
| Reagent or resource | | |
| Oil red O | Sigma-Aldrich | Cat#: O1391 |
| TRIzol Reagent | Thermo Fisher Scientific | Cat#: 15596026 |
| RIPA Buffer | Boston BioProducts | Cat#: BP-115 |
| BCA assay kit | Thermo Fisher Scientific | Cat#: 23227 |
| West Femto Maximum Sensitivity Substrate | Thermo Fisher Scientific | Cat#: 34095 |
| 0.05% Trypsin 0.53 mM EDTA | Corning | Cat#: MT25052CI |
| Critical Commercial Assays | | |
| Fixation/Permeabilization Solution Kit | BD Biosciences | Cat#: 554714 |
| High Capacity cDNA Reverse Transcription Kit | Thermo Fisher Scientific | Cat#: 4368813 |
| 2X Universal SYBR Green Fast qPCR mix | ABclonal | Cat#: RK21203 |
| Experimental Models: Organisms/Strains | | |
| Mouse: *TBX18-Cre*$^{ERT2}$ | Cuimaraes-Camboa et al., 2017 | #031520 |

| | | |
|---|---|---|
| Mouse: *Ucp1*-Cre$^{ERT2}$ | Rosenwald et al., 2013 | N/A |
| Mouse: *PDGFRβ-Cre$^{ERT2}$* | Henar Cuervo et al., 2017 | #030201 |
| Mouse: *Rbpj*$^{fl/fl}$ | Han et al., 2002 | N/A |
| Mouse: *Rosa26R*$^{RFP}$ | Jackson laboratory | #007914 |
| Mouse: *Rosa26R$^{DTA}$* | Jackson laboratory | #006331 |
| Oligonucleotides | | |
| A full list of qPCR primers, see Table S1 | This paper | N/A |
| Software and Algorithms | | |
| Word | Microsoft | N/A |
| Excel | Microsoft | N/A |
| ImageJ | NIH | https://imagej.nih.gov/ij/ |
| Prism | GraphPad Software | Graphpad Software |
| Other | | |
| Zeiss LSM8800 Confocal Microscope | Zeiss | N/A |
| Leica DMi8 microscope | Leica | N/A |
| Leica M205 FA microscope | Leica | N/A |
