## Supplementary table 1 for "The Notch-Pdgfrβ axis suppresses brown adipocyte progenitor differentiation in early postnatal mice"

**Table S1, Related to STAR Methods.**

**Sequences of primers.**

| Gene | Species | Forward primer (5'-3') | Reverse primer (5'-3') |
|---|---|---|---|
| *Actin* | Mouse | CCACTGGCATCGTGATGGACTCC | GCCGTGGTGGTGAAGCTGTAGC |
| *Pdgfrβ* | Mouse | TGTGCAGTTGCCTTACGACT | CAGGTGGGGTCCAAGATGAC |
| *Pdgfd* | Mouse | CTAGGTCGCTGAGCCCAAAT | CATCTCTCCTGAGGTTGGCA |
| *Rbpj* | Mouse | GAATTTCCACGCCAGTTCAC | ATACAGGGTCGTCTGCATCC |
| *Notch2* | Mouse | GTGACCATCTTTCGGAGGCA | GCGAGAATGTCTGGGCGATA |
| *Notch3* | Mouse | TCAGAATGTCCAGAGGCCAAG | GTGAAGCCATCAGGCCCTC |
| *Hes1* | Mouse | GCAGACATTCTGGAAATGACTGTGA | GAGTGCGCACCTCGGTGTTA |
| *Hey1* | Mouse | CATGAAGAGAGCTCACCCAGA | CGCCGAACTCAAGTTTCC |
| *Cd36* | Mouse | TTTGGAGTGGTAGTAAAAAGGGC | TGACATCAGGGACTCAGAGTAG |
| *Gapdh* | Mouse | TGGATTTGGACGCATTGGTC | TTTGCACTGGTACGTGTTGAT |
| *Ldha* | Mouse | TGTCTCCAGCAAAGACTACTGT | GACTGTACTTGACAATGTTGGGA |
| *Pfkp* | Mouse | CGCCTATCCGAAGTACCTGGA | CCCCGTGTAGATTCCCATGC |
| *Cpt1a* | Mouse | CATCACCCCAACCCATATTC | CGTGTTGGATGGTGTCTGTC |
| *Cpt1b* | Mouse | TGTCATGGCAACAGTTGGTT | GACTCCGGTGGAGAAGATGA |
| *Lpl* | Mouse | GGGAGTTTGGCTCCAGAGTTT | TGTGTCTTCAGGGGTCCTTAG |
| *Hsl* | Mouse | ACGCTACACAAAGGCTGCTT | TCGTTGCGTTTGTAGTGCTC |
| *ATGL* | Mouse | TTCACCATCCGCTTGTTGGAG | AGATGGTCACCCAATTTCCTC |
| *Glut4* | Mouse | CTCATGGGCCTAGCCAATG | GGGCGATTTCTCCCACATAC |
| *Fasn* | Mouse | TTGACGGCTCACACACCTAC | CGATCTTCCAGGCTCTTCAG |
| *Eno1* | Mouse | TGCGTCCACTGGCATCTAC | CAGAGCAGGCGCAATAGTTTTA |
| *Hk2* | Mouse | TGATCGCCTGCTTATTCACGG | AACCGCCTAGAAATCTCCAGA |
| *Pkm2* | Mouse | GCCGCCTGGACATTGACTC | CCATGAGAGAAATTCAGCCGAG |
| *Leptin* | Mouse | AAGACCATTGTCACCAGGATCAA | GGATACCGACTGCGTGTGTG |
| *Cox8b* | Mouse | GAACCATGAAGCCAACGACT | GGCGTAGCTCCTCAAACAAC |

| | | | |
|---|---|---|---|
| *Elovl3* | Mouse | TTCTCACGCGGGTTAAAAATGG | GCGAAGTTCACAGTGGTTCC |
| *Dio2* | Mouse | ACACTGGAATTGGGAGCATC | ATGCTGACCTCAGAAGGGCT |
| *Prdm16* | Mouse | ACACGCCAGTTCTCCAACCTGT | TGCTTGTTGAGGGAGGAGGTA |
| *Pparg2* | Mouse | GCATGGTGCCTTCGCTGA | TGGCATCTCTGTGTCAACCATG |
| *Ppargc1a* | Mouse | CCGATCACCATATTCCAGGT | GTGTGCGGTGTCTGTAGTGG |
| *Cidea* | Mouse | TCTGCAATCCCATGAATGTC | CAGTGATTTAAGAGACGCGG |
| *Ucp1* | Mouse | CACCTTCCCGCTGGACACT | CCCTAGGACACCTTTATACCTAATGG |
| *aP2* | Mouse | CTGGGCGTGGAATTCGAT | GCTCTTCACCTTCCTGTCGTCT |
| *Leptin* | Mouse | AAGACCATTGTCACCAGGATCAA | GGATACCGACTGCGTGTGTG |
| *Rfp* | Mouse | GCTTCAAGGTGCGCATGGAG | CGGTGTTGTGGCCCTCGTAG |
